# Lipid bilayers determine allostery but not intrinsic affinity of cAMP binding to pacemaker channels

**DOI:** 10.1101/2024.12.23.630133

**Authors:** Vinay Idikuda, Susovan Roy Chowdhury, Yongchang Chang, Qian Ren, Huan Bao, Randall H. Goldsmith, Baron Chanda

**Affiliations:** Department of Anesthesiology, Washington University in St. Louis; Saint Louis, MO 63110; Department of Chemistry, University of Wisconsin-Madison; Madison, WI 53706; Department of Biochemistry and Molecular Biophysics, Washington University School of Medicine; Saint Louis, MO, USA; Department of Neuroscience, Washington University School of Medicine; Saint Louis, MO, USA; Department of Molecular Physiology and Biological Physics, University of Virginia; 480 Ray C Hunt Drive, Charlottesville, 22903, Virginia, USA

**Keywords:** Single-molecule-spectroscopy, Co-operativity, lipids, ion-channels, HCN channels

## Abstract

Cyclic adenosine monophosphate (cAMP), a second messenger, binds to hyperpolarization and cyclic nucleotide-gated (HCN) ion channels and regulates the automaticity of pacemaking activity. While cellular studies suggest that cAMP binding to HCN channels exhibits unusual cooperativity, recent findings using purified detergent-solubilized channels indicate independent binding to each subunit. This discrepancy raises the question of whether the lipid environment or endogenous cellular cofactors influence cAMP-dependent gating. To address this, we reconstituted purified human HCN channels in nanodiscs and resolved cAMP binding energetics at single-molecule resolution using nanophotonic waveguides. Our measurements reveal that, in contrast to detergent-solubilized channels, cAMP binds cooperatively to HCN channels reconstituted in a variety of lipid nanodiscs. Remarkably, the presence of lipid bilayer promotes ligand-binding allostery but not intrinsic binding affinity. To explore the molecular basis of bilayer-induced allostery, we determine the cryo-EM structure of HCN1 in soy polar lipid nanodiscs at a nominal resolution of 3.77 Å resolution. Although the overall architecture is conserved, the average interfacial distance between the transmembrane domain and C-terminal domain of neighboring subunits are shorter in lipid nanodiscs. These findings indicate that the lipid bilayer regulates the function of pacemaker ion channels by enhancing inter-subunit interactions and underscore the fundamental role of membranes in amplifying the gating sensitivity of ion channels by promoting long-range cooperative interactions.

**Significance Statement:** Lipid membranes are essential for the structure and function of membrane proteins, including ion channels. Lipid mimetics, such as non-ionic detergents, are widely used as surrogates for the membrane environment in structural and biophysical studies. Here, we demonstrate that while the overall structure of the pacemaker ion channel remains similar, lipid membranes—unlike detergents—promote cooperative ligand-binding transitions by modifying interactions at intersubunit interfaces. These findings provide new insights into the mechanism of ion channel regulation by lipid membranes.

## Introduction

HCN, or pacemaker channels, are a distinct subclass of voltage-gated ion channels that are activated by membrane hyperpolarization and modulated by the intracellular second messenger cAMP(1-5). By integrating electrical and chemical signals, HCN channels play a critical role in regulating the rhythmic firing of cardiac and neuronal action potentials(6, 7). Each subunit of a tetrameric HCN channel contains a single cAMP binding site in its intracellular cyclic nucleotide-binding domain (CNBD). The binding of cAMP allosterically modulates channel gating, enhancing macroscopic current and accelerating activation kinetics(8).

Despite the essential role of cAMP in pacemaking, the mechanisms underlying cAMP-dependent gating of HCN channels remain controversial. Electrophysiology and patch-clamp fluorometry studies suggest that HCN channels function as dimer-of-dimers, with higher binding affinities for the first and third ligation states compared to the unbound or doubly-bound states(9-15). In contrast, cAMP binding to isolated CNBDs promotes tetramerization rather than dimerization suggesting a canonical positive cooperativity(16). This inconsistency is not surprising because ensemble measurements lack the ability to discriminate between various models due to issues of parameter identifiability(17). By measuring the distributions of individual ligation states, single-molecule measurements can unambiguously settle this question(17, 18). However, detecting individual ligand binding events at concentrations above 10 nM is difficult due to the increased background from free fluorescent ligands(19-21). Recently, White et al. utilized zero-mode waveguides (ZMWs) to demonstrate that purified HCN channels bind all four cAMP molecules independently(22, 23). ZMWs reduce the excitation volume by three orders of magnitude compared to conventional TIRF microscopy, enabling the study of fluorescent ligand binding dynamics at micromolar concentrations(24-29).

Although isolated HCN channels bind cAMP molecules independently, it remains unclear whether extrinsic factors, such as surrounding lipids or other cofactors modulate ligand binding in cellular environments. High-resolution structures of detergent-solubilized HCN channels reveal bound lipid densities consistent with previous functional studies(30, 31). However, a vast majority of the surrounding lipids, known as solvating lipids, interact transiently with integral membrane proteins(32-34). They are replaced by detergents upon solubilization and are not resolved in the high-resolution structures. While the influence of specific lipids or physical properties of the bilayer on channel structure and function has been extensively studied(31, 35-39), the role of these solvating lipids vis-a-vis other membrane mimetics is less understood. Since most structures of membrane signaling proteins, such as ion channels and GPCRs, are solved in detergents(40, 41), understanding how these environments differ is crucial for interpreting functional mechanisms accurately and guiding future structural studies.

To address these questions, we used HCN channels which are uniquely suited for studying ligand-dependent gating energetics at single-molecule resolution in both detergents and lipid environment(22, 42). Here, we combined single-molecule ligand-binding studies with single-particle cryo-EM to examine HCN1 and HCN4 channels in lipid nanodiscs of varied compositions and compared them to detergents widely used for structural studies. Our single-molecule studies reveal that, in contrast to detergents, lipid bilayers cause cAMP to bind cooperatively without altering the intrinsic binding affinity. Cryo-EM analysis further shows that structural rearrangements in lipid nanodiscs promote intersubunit interactions. These findings provide new insights into how lipid bilayers modulate the quaternary structure and influence the allosteric regulation of pacemaker ion channels.

## Results

### HCN1 reconstituted in soy lipid nanodiscs exhibits increased ligand binding activity

To investigate whether lipid bilayers influence the ligand-binding dynamics of HCN1 channels, we isolated and purified heterologously expressed HCN1 channels using glyco-diosgenin (GDN) detergent. A portion of these purified channels was reconstituted into soy polar lipid nanodiscs by mixing the HCN1 protein with the scaffold protein MSP2N2 and phospholipids in a 1:2.5:200 ratio (Fig. 1A, Table 1). Reconstitution was validated through size exclusion chromatography and SDS-PAGE analysis (Fig. 1B), with tetrameric assembly confirmed via GFP photobleaching (Fig. S2).

**Table 1.** Molar Ratios of HCN1, scaffold proteins and lipids for various nanodisc reconstitution assays.

| Protein | Scaffolding protein | Lipids | Molar Ratio (Protein: MSP: Lipid) |
| --- | --- | --- | --- |
| HCN1 | MSP2N2 | Soy Polar lipids | 1:2.5:200 |
| HCN1 | MSP2N2 | POPC/POPE | 1:2.5:280 |
| HCN1 | MSP2N2 | Brain Polar lipids | 1:2.5:200 |
| HCN1 | spNW11 | Brain Polar lipids | 1:2.5:95 |
| HCN1 | spNW30 | Brain Polar lipids | 1:2.5:600 |
| HCN4 | MSP2N2 | Soy Polar lipids | 1:2.5:200 |

**Fig. 1.**
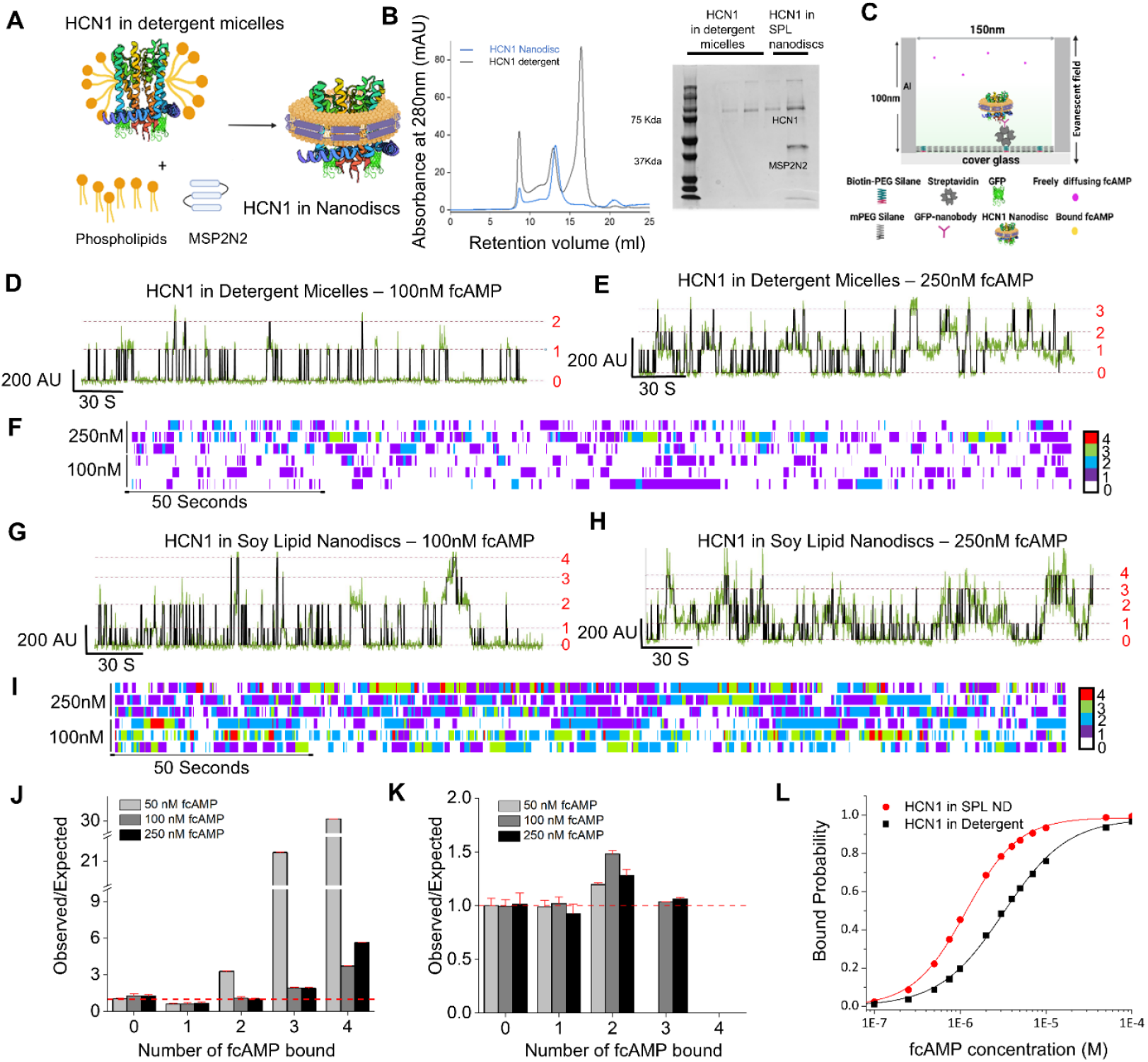
Ligand-binding activity in nanodisc-reconstituted and detergent-solubilized HCN channels. **(A)** Schematic of HCN1 reconstitution into lipid nanodiscs using MSP2N2 as scaffold protein (1:2.5:200 ratio of HCN1, MSP2N2, and soy polar lipids). **(B)** (left) Size exclusion chromatography profile of HCN1 purified in detergent (blue) and HCN1 purified in detergent and reconstituted into soy polar lipid nanodiscs with MSP2N2 (black). (right) Coomassie blue stained SDS-PAGE gel showing various SEC fractions of HCN1 at ∼75Kda before reconstitution (lanes 2-4) and both HCN1 and MSP bands after reconstitution (lane 5) **(C)** Cartoon (not to scale) of single-molecule pulldown of HCN1 onto a ZMW. ZMWs passivated with mPEG silane doped with biotin-PEG-silane and coated with streptavidin. Biotinylated GFP pulled down GFP-tagged HCN1. **(D, E)** Representative FT traces of 100 nM fcAMP (**D**) and 250 nM fcAMP (**E**) binding to HCN1 in a detergent micelle. Dashed lines, ligated state of the channel. **(F)** Heatmap of three independent FT traces of 100 nM and 250 nM fcAMP binding to HCN1 in detergent micelles. **(G, H)** Representative FT traces of 100 nM and 250 nM fcAMP binding to HCN1 in soy polar lipid nanodiscs. **(I)** Heatmap of three independent FT traces of 100 nM and 250 nM fcAMP binding to HCN1 in soy polar lipid nanodiscs. **(J, K)** Ratios of observed (P(k)_obs_) and expected (P(k)_exp_) values of liganded state occupancies from binomial fits across all trajectories for HCN1 in soy polar lipid nanodiscs (**J**) or detergent micelles (**K**) at different fcAMP concentrations. Red dashed line, concordance between observed and expected values; error bars (red), standard deviation. Soy polar lipid nanodiscs, N = 945 molecules; total observation time 112.24 hours; total events 234859 (table S1). Detergent, N = 579 molecules; total observation time 50.5 hours; total events 106542 (table S2). **(L)** Extrapolated binding curve for HCN1 in soy polar lipid nanodiscs (black) and detergent (red) generated by calculating predicted bound probability from experimentally obtained microscopic equilibrium rate constants and substituting them in Adair’s equation for four binding sites(46).

We measured ligand binding by immobilizing purified channels—either in GDN micelles or lipid nanodiscs— onto zero-mode waveguides (ZMWs) and introducing the fluorescent ligand fcAMP (DY-547-cAMP) (Fig. 1C). To ensure single-occupancy of ZMWs, protein concentration was adjusted based on Poisson dilution statistics as previously described(43). Ligand binding events were measured across a range of fcAMP concentrations using micromirror total internal reflection fluorescence (TIRF) microscopy(44). Fluorescence intensity-time (FT) traces were analyzed using autoDISC(23, 45) to identify bound and unbound states (Fig. 1D-I, Fig. S3). Control experiments confirmed the absence of nonspecific binding, as evidenced by no detectable ligand binding in the absence of HCN channels and by the competitive displacement of fcAMP with excess non-fluorescent cAMP at the end of each experiment (Fig. S4).

Our results revealed that channels in nanodiscs exhibited a higher frequency of multi-ligand bound states (three or four fcAMP molecules bound) compared to the singly bound state, particularly at lower fcAMP concentrations (Fig. 1D-E, Fig. 1G-H). To quantify this effect, we normalized the time spent by fcAMP in each ligation state (state occupancy) to the expected value from a binomial distribution under the assumption of independent and identical binding sites. When ligand binding is independent, the ratio of observed to expected occupancies is 1. For HCN1 channels in nanodiscs, occupancy probabilities deviated significantly from binomial predictions, especially in high-occupancy states across all concentrations tested (Fig. 1J, Fig. S7B, Table S10). For instance, at 50 nM fcAMP, the fourth bound state showed a deviation exceeding 25-fold. This effect diminished at higher fcAMP concentrations (2-5-fold deviation) as we reached saturation. In contrast, channels in detergent micelles displayed binding distributions consistent with independent binding (Fig. 1K, Fig. S7A, Table S11), consistent with previous observations(22).

Using a sequential binding model, we calculated bound probabilities for each ligation state at various concentrations and generated ensemble binding curves from the single-molecule data. A bound probability of 1.0 indicates complete occupancy of all binding sites by fcAMP. From these probabilities, we determined the microscopic equilibrium constants for each transition (see Methods). Using the Adair equation(46), we constructed binding curves (Fig. 1L). HCN1 channels in nanodiscs showed a three-fold increase in macroscopic binding affinity compared to those in detergent micelles (Fig. 1L, black vs. red).

Overall, these results demonstrate that fcAMP binds to HCN1 channels in nanodiscs non-independently, in contrast to those in detergent micelles.

### Brain and neutral lipids both also promote occupancy of higher-order ligation states

We wondered whether the cooperativity observed in HCN1 channels reconstituted in soy polar lipids was influenced by specific lipid species. Phospholipids are key regulators of many ion channels in biological membranes(47-52), and recent studies(31) show that negatively charged phospholipids modulate HCN channel activity. To test this, we compared two lipid conditions: a native lipid composition (porcine brain polar lipid extract) and a more straightforward neutral lipid system (POPC/POPE in a 3:1 ratio). HCN channels exhibited robust binding activity at all concentrations in both brain polar lipid and POPC/POPE nanodiscs (Fig. 2A and B, Fig. S5).

**Fig. 2.**
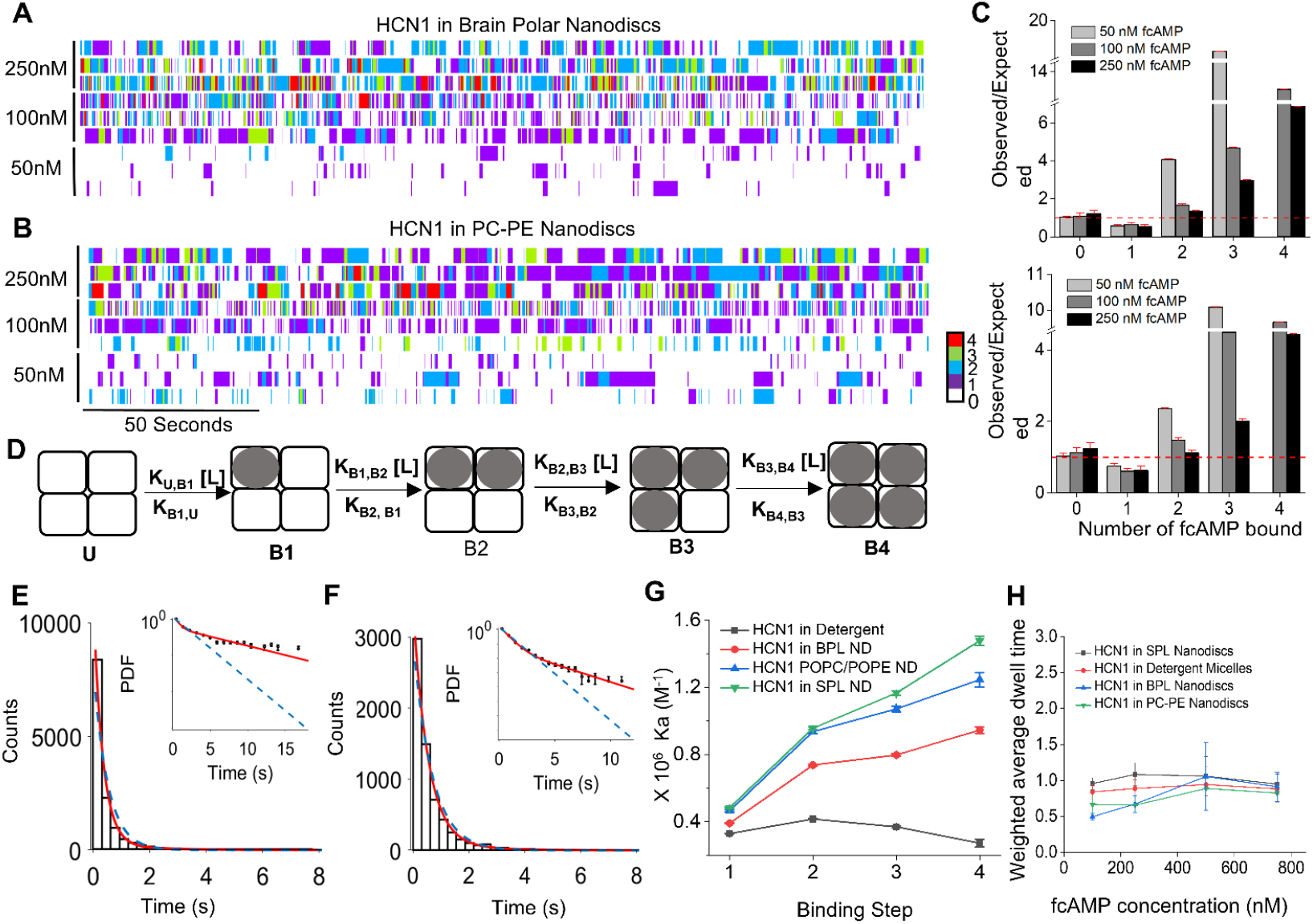
Lipid bilayer alter microscopic binding affinities for higher-order ligation states. **(A, B)** Heatmap of three independent FT traces of 50 nM, 100 nM, and 250 nM fcAMP binding to HCN1 reconstituted in (**A**) porcine brain polar lipid nanodiscs and (**B**) POPC/POPE lipid nanodiscs. **(C)** Ratios of P(k)_obs_ and P(k)_exp_ for liganded state occupancies from binomial fits across all trajectories for HCN1 in brain polar lipid nanodiscs (top) and a 3:1 ratio of POPC/POPE (bottom) at different fcAMP concentrations. Brain polar lipid nanodiscs, N = 898 molecules; total observation time = 90.24 hours; total events = 238280. POPC/POPE nanodiscs, N = 372 molecules; total observation time = 51.2 hours, total events = 141600. See tables S3 and S4. (**D**) Schematic of sequential model for ligand binding. White squares unbound (U); squares with grey circles, bound (B) ligand. **(E, F)** Dwell time distributions of isolated UBU events for HCN1 in (**E**) brain polar lipid nanodiscs and (**F**) POPE/POPE nanodiscs at 100 nM fcAMP. Distributions overlaid with maximum likelihood estimates for monoexponential (blue dashed) and bi-exponential (red) distributions. Error bars denote standard deviation. Individual time constants and amplitudes in table S21, S22. **(G)** Microscopic equilibrium association constants (Ka) for each ligation state of HCN1 in Soy polar lipid nanodiscs, POPC/POPE nanodiscs, brain polar lipid nanodiscs, and detergent micelles. Parameters obtained by global fitting to all events across all concentrations for each condition. (**H)** Biexponential weighted dwell time constant of the first bound state at various fcAMP concentrations obtained by fitting UBU dwell times with a biexponential function.

Using idealized FT traces, we determined state occupancies for each ligation state across the concentration range and normalized them to expected values based on an independent binding model. In both brain polar lipids (Fig. 2C, top) and POPC/POPE (Fig. 2C, bottom), HCN1 state occupancies deviated from the expected values, mirroring trends observed in soy polar lipids showing non-independent binding.

Next, we calculated the bound open probabilities and estimated the microscopic rate constants using fits to an unconstrained sequential binding model as described previously (see Methods)(Fig. 2D). The results showed a monotonic but non-linear increase in microscopic equilibrium constants across all lipid nanodisc types (Fig. 2G, red, blue, green, and tables S28, S29, S30). These findings demonstrate that fcAMP binding exhibits positive cooperativity in lipid environments, regardless of lipid origin or charge, but not in detergent-solubilized channels.

Next, we examined whether lipid-induced cooperativity arises from changes in agonist binding affinity or subsequent allosteric transitions. We analyzed unbound-bound-unbound (UBU) dwell time distributions for singly bound states, avoiding any events that only show the binding of new ligands or the unbinding of previously bound ones(22). UBU dwell times correspond to the binding of the first ligand, and the analysis of this bound dwell time gives us an estimate of intrinsic ligand binding affinity without confounding effects of allosteric regulation.

Consistent with our prior work, maximum likelihood estimations of UBU dwell time distributions for HCN1 were well described by two exponentials at all fcAMP concentrations (Fig. 2E and 2F, Fig. S9A and B, Fig. S10A and B, and tables S19, S20, S21, S22). This revealed that each subunit undergoes isomerization to a secondary conformation or flip state(22). Importantly, these biexponential fits were consistent across detergent and nanodisc samples, and their weighted time constants were similar in all conditions (Fig. 2H and tables S19, S20, S21, S22). These results suggest that the lipid environment does not alter the intrinsic affinity of fcAMP for its binding site or the time spent in the initial bound state. Instead, lipids influence downstream allosteric transitions and the energetics of higher-order ligand-binding occupancies.

### Cooperative binding of cAMP is not isoform-specific or dependent on nanodisc size

Recent findings have demonstrated that the structure and function of certain membrane proteins depend on nanodisc size(53-55). While MSP2N2 nanodiscs have an average diameter of 15 nm, their size distribution is broad and influenced by the reconstituted protein. To examine whether nanodisc size impacts the cooperativity of ligand binding, we used circular nanodiscs with uniform sizes across a wide temperature range and minimal dependence on the reconstituted protein(56, 57). Spy-catcher and spy-tag MSPs were purified (Fig. S1C and S1, F-I) to generate circular nanodiscs of 11 nm and 30 nm using soy polar lipids. Heatmap representations of FT traces revealed strong ligand-binding activity for both nanodisc types (Fig. 3A and Fig. S6). Even at the lowest tested concentration (50 nM fcAMP), higher-order ligation states were observed, contrasting with detergent-solubilized samples.

**Fig. 3.**
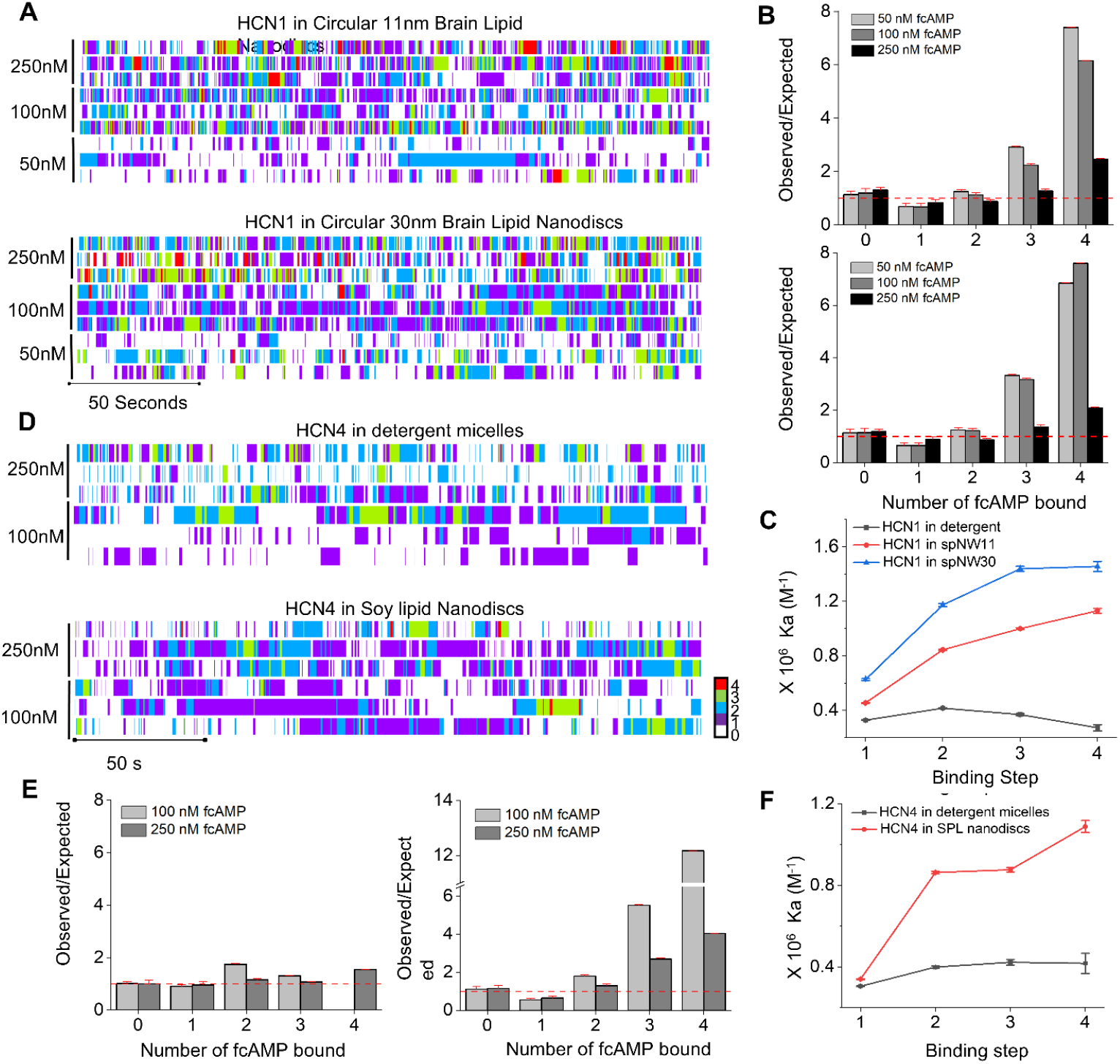
Ligand-binding cooperativity in different-sized circular nanodiscs and HCN4 isoform. **(A)** Heatmap of three independent FT traces of 100 nM, 250 nM, and 750 nM fcAMP binding to HCN1 reconstituted in 11 nm (spMSP1D1, top) and 30 nm (spNW30, bottom) circular nanodiscs of brain polar lipids. **(B)** Ratios of P(k)_obs_ and P(k)_exp_ for liganded state occupancies from binomial fits across all trajectories for HCN1 reconstituted into 11 nm (top) and 30 nm (bottom) circular nanodiscs of brain polar lipids at 50 nM, 100 nM and 250 nM fcAMP. 11 nm circular nanodiscs, N = 743 molecules; total observation time 122.3 hours; total events 379403. 30 nm circular nanodiscs, N = 461 molecules; total observation time 76.4 hours; total events 198934. See tables S5 and S6. **(C)** Heatmap of three independent FT traces of 100 nM and 250 nM fcAMP binding to HCN4 in GDN detergent micelles (top) and soy polar lipid nanodiscs (bottom). **(D)** Ratios of P(k)_obs_ and P(k)_exp_ for liganded state occupancies from binomial fits across all trajectories for HCN4 reconstituted in detergent micelles (top) and soy polar lipid nanodiscs (bottom) at 100 nM and 250 nM fcAMP. Detergent micelles, N = 517 molecules; total observation time 43.08 hours; total events 43982. Soy polar lipid nanodiscs, N = 498 molecules; total observation time 58 hours; total events 97429. See tables S7 and S8. **(E)** Ka values for all tested conditions showing positive cooperativity for HCN1 in both 11 nm and 30 nm nanodiscs, and independent binding in detergent micelles. Parameters obtained by fitting an optimized unconstrained sequential model to all events across all concentrations. **(F)** Ka values for all tested conditions showing positive cooperativity for HCN4 in soy lipid nanodiscs and independent binding in detergent micelles. Parameters fit to all events across all concentrations.

Plots comparing observed and expected state occupancies for unbound and various bound states deviated from the binomial distribution, indicating non-independent binding (Fig. 3B, Fig. S8, B and C, Fig. S10, C and D, and tables S23, S24). Using the unconstrained sequential binding model in Fig. 2D, we extracted microscopic equilibrium constants for each of the four binding steps from idealized FT traces (Fig. 3C). Consistent with data from MSP2N2 nanodiscs, HCN1 reconstituted in both 11 and 30 nm circular nanodiscs exhibited positive cooperativity. Notably, the strongest cooperativity was observed in 30 nm nanodiscs, where the microscopic equilibrium constants for the third and fourth ligation states were nearly identical (Fig. 3C, blue). This finding aligns with functional data indicating that 60% cAMP occupancy is sufficient for maximal channel activation(11).

To determine whether this effect is generalizable to other HCN channels, we expressed and purified a C-terminal truncated, GFP-tagged human ortholog of HCN4 (Fig. S1, J, K, L), which is approximately four times more sensitive to cAMP than HCN1(58). These channels were solubilized in GDN (Glyco-diosgenin), with a portion reconstituted in soy polar lipid nanodiscs using MSP2N2 as the MSP. Heatmap representations of FT traces showed that HCN4 ligand-binding activity in detergent micelles was lower than in soy polar lipids (Fig. 3D). In detergent micelles, the ratios of observed to expected occupancies for various ligation states were close to one, consistent with independent ligand-binding events (Fig. 3E, left). By contrast, observed occupancies in soy polar lipids deviated from independence (Fig. 3E, right), resembling HCN1 channels reconstituted in nanodiscs (Fig. S12, A and B, table S16, S17).

Similar to HCN1 and HCN2(22), UBU dwell-time analysis of HCN4 revealed that the channel undergoes secondary isomerization into a flip state (Fig. S12, C and D, and tables S16, S17, S26, and S27). Importantly, extracted microscopic equilibrium constants for each ligation step indicated that binding of the second and subsequent ligands is more favorable in channels reconstituted in nanodiscs than in detergent-solubilized channels (Fig. 3F, tables S35 and S36). Thus, like HCN1 isoform, ligand binding to HCN4 channel reconstituted in lipid bilayers exhibits positive cooperativity, unlike those solubilized in detergent micelles.

### HCN1 channels in native membranes also bind to cAMP cooperatively

Reconstituting membrane proteins in nanodiscs requires solubilizing both the target protein and lipids in detergent—in this case, GDN and CHAPS, respectively. Although we use BioBeads to remove most, if not all, detergent, we wondered whether exposure to detergent during purification might irreversibly alter the channel gating energetics. To address this question, we employed detergent-free nanodiscs (DeFrNDs), which circumvent the need for detergent-mediated solubilization and reconstitution steps(59). These nanodiscs are a novel class of engineered membrane scaffold peptides derived from apolipoprotein A that facilitate the direct extraction of cell membranes as nanoscale membrane discoids (Fig. 4A). The HCN1-encapsulated discoids were further isolated by affinity purification.

**Fig. 4.**
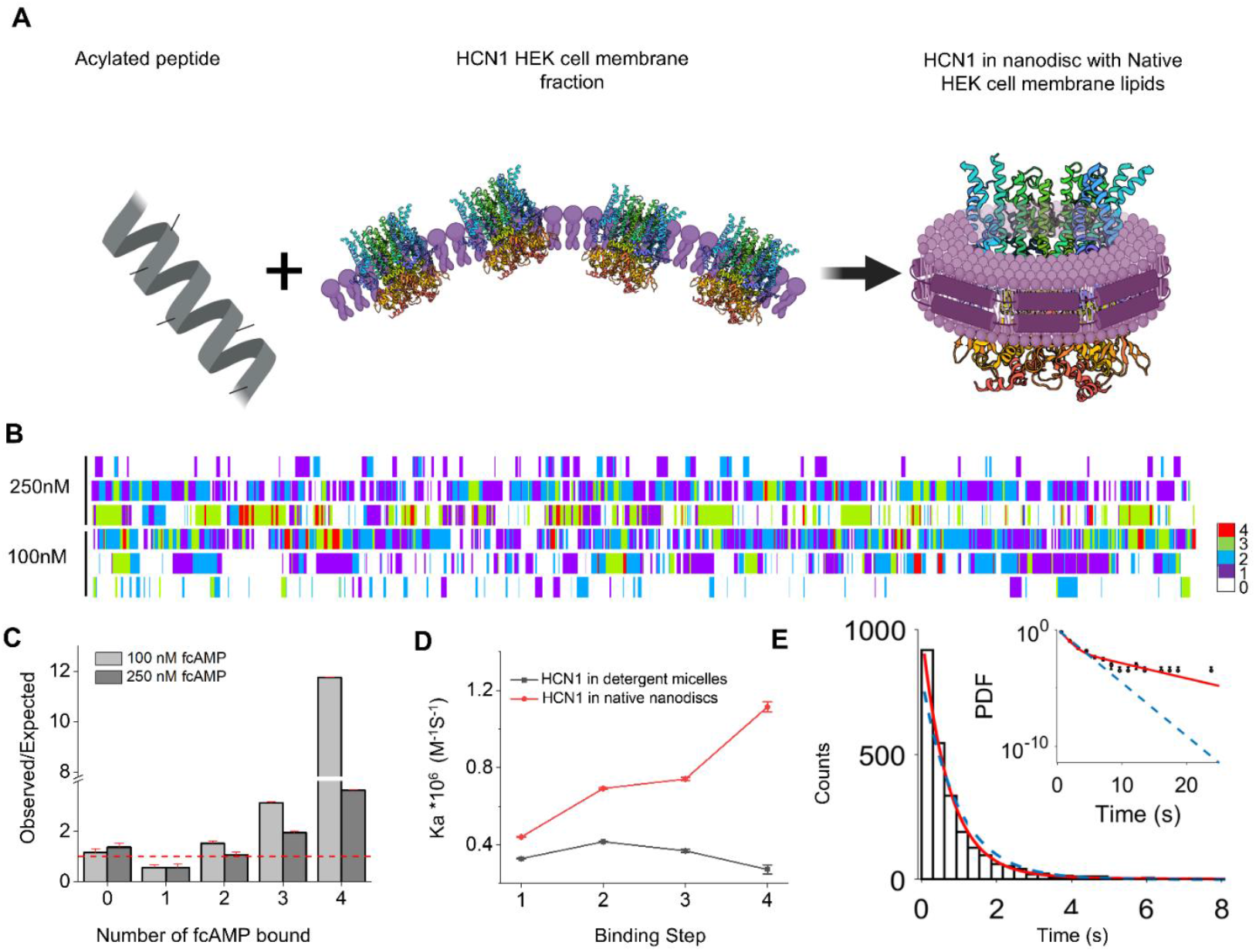
HCN channels purified in native nanodiscs exhibit show cooperative fcAMP binding. **(A)** Schematic of HCN1 reconstitution into native HEK cell nanodiscs using acylated peptides as a MSP. **(B)**Ratios of P(k)_obs_ and P(k)_exp_ for liganded state occupancies from binomial fits across all trajectories for HCN1 in native HEK membrane lipid nanodiscs at 100nM and 250nM fcAMP. **(C)** Heatmap of three independent FT traces of 100nM and 250nM fcAMP binding to HCN1 purified in native HEK cell nanodiscs. **(D)** Ka values for all tested conditions showing positive cooperativity for HCN1 in native lipid nanodiscs and independent binding in detergent micelles. Parameters fit to all events across all concentrations. Native HEK cell nanodiscs, N = 671 molecules. See table S9. **(E, F)** Dwell time distributions of isolated B1 events for HCN1 in native HEK cell nanodiscs at 100nM (**E**) fcAMP. Distributions overlaid with maximum likelihood estimates for monoexponential (blue dashed) and bi-exponential (red) distributions. Error bars are for a binomial distribution. Individual time constants and amplitudes in table S25.

HCN1 channels extracted into DeFrNDs directly from HEK cells exhibited robust ligand-binding activity, as evidenced by a high binding activity and substantial occupancy of higher ligation states at low concentrations (Fig. 4B, Fig. S13A). Similar to HCN channels in detergents and nanodiscs, HCN1 channels in DeFrNDs showed UBU dwell time distributions with two time constants, indicating isomerization occurs after ligand binding (Fig. 4E, table S34). Furthermore, the observed state occupancies deviated significantly from expected binomial distributions (Fig. 4C, table S18), and microscopic equilibrium constants derived from globally fitted FT traces at all concentrations showed that the higher ligation states were favored (Fig. 4D, Fig. S13B). Thus, HCN1 channels in DeFrNDs exhibit positive cooperativity comparable to that observed in reconstituted nanodiscs following detergent solubilization.

### Structural analysis of detergent and nanodisc reconstituted HCN channels

To determine whether lipid bilayers and detergent micelles differentially affect the structure of HCN channels, we reconstituted human HCN1 in circular nanodiscs and compared the single-particle cryo-EM structure to those obtained with detergent(60). Due to strong orientation bias, we collected 30° tilt series data and combined them with a small fraction of particles from a non-tilted sample containing 0.2 mM fluorinated fos-choline to further reduce orientation preference caused by protein accumulation at the air-water interface.

The resulting map closely resembles the closed channel conformation in detergent micelles (Fig. 5A, figs. S14 and S15). Using the closed-state structure as the starting model, we built a structural model for hHCN1-nanodisc. Supplemental videos show the HCN1 structure transitioning from detergent to nanodisc, highlighting primary changes in inter-subunit interfaces. These include shortening the distances between the S4-S5 loop and the C-linker, as well as between the HCN domain and the CNBD (movies S1 and S2). This is further illustrated in Fig. 5C, which superposes the hHCN1-nanodisc structure onto the hHCN1-detergent structure. Upward movement of the C-linker A’ and B’ and the CNBD domain in the nanodisc model resulted in an average decrease of ∼1Å between these regions compared to the detergent model.

**Fig. 5.**
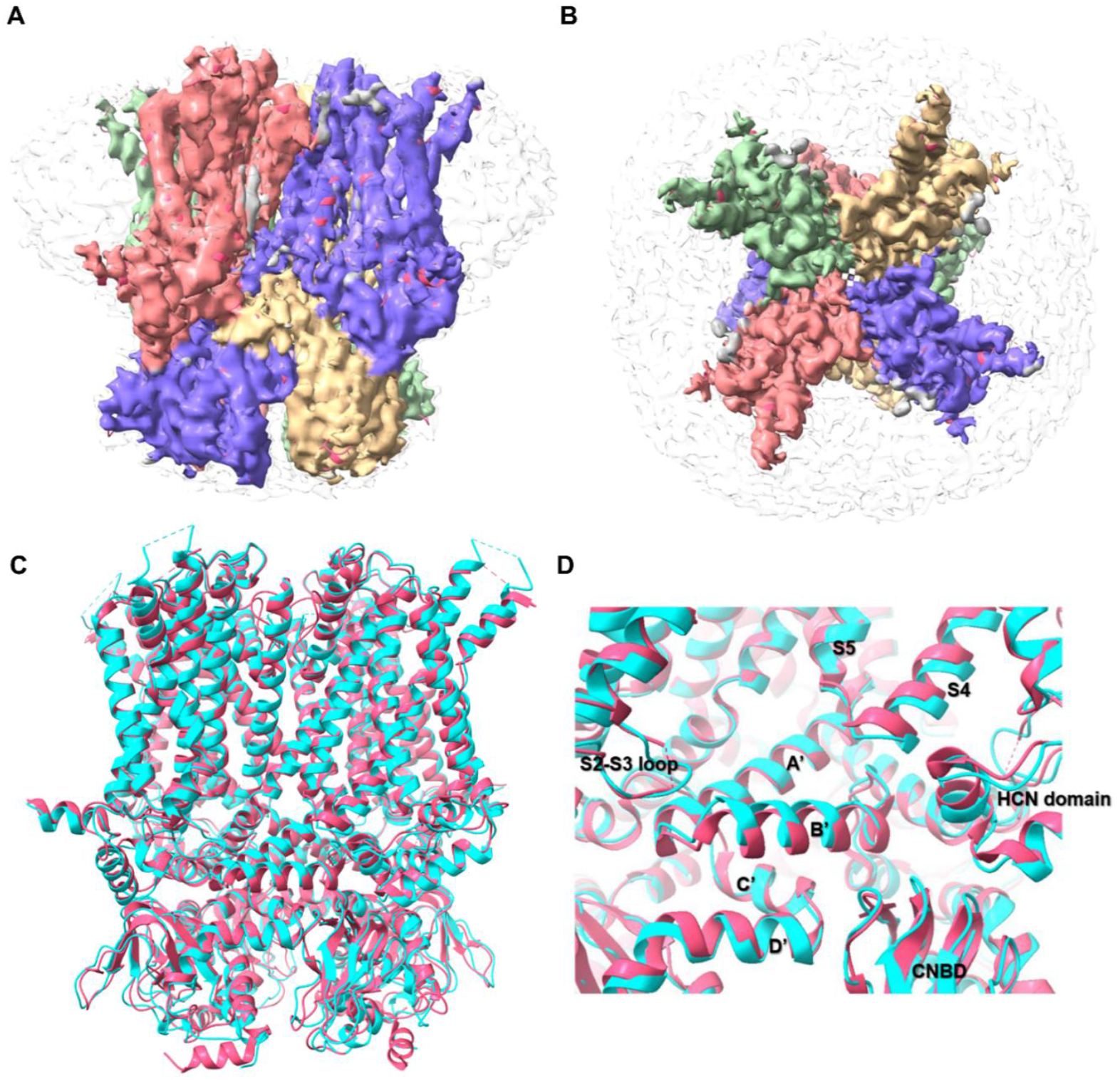
Cryo-EM structure of hHCN1 in soy polar lipid nanodisc. **(A**) Side-view of hHCN1 cryo-EM map colored with the superimposed model subunits. **(B)** Superimposed models of hHCN1-nanodisc (cyan) and hHCN1-detergent (PDB: 5u6o, red) zoomed in to the region with inter-subunit interactions. **(C)** Comparison of hHCN1-nanodisc and hHCN1-detergent at the interface between S4-S5/HCN and C-terminal domain (including C-linker and CNBD). Left: structural models with ribbon presentation; Right: structural models with surface presentation. Note that a small upward movement of A’-B’ of the C-linker and CNBD and a small downward movement of HCN domain and S4-S5 loop. **(D)** Comparison of hHCN1-nanodisc and hHCN1-detergent at the interface between C’-D’ of the C-linker and C-terminal domain (including A’-B’ of the C-linker on the upper left and CNBD on the upper right).

Fig. 5D shows zoomed-in views of two inter-subunit interfaces where structural deviations between the two conformations are observed. Narrowing of the cleft at the interface comprising the S4-S5 loop, HCN domain, and CNBD from the neighboring subunit is evident in the surface model of the nanodisc structure (Fig. 5C). We observe a similar narrowing of the cleft in the recently published structure of human HCN1 in PC:PE:PS lipid nanodiscs(61) (PDB: 8UC8) (Fig. S16). Fig. 5D highlights additional changes in the inter-subunit interface between the CNBD and C-linker region. In the nanodisc structure, the C’ and D’ helices of the C-linker are further from the A’ and B’ helices of the neighboring C-terminal domain but closer to the β-roll of the CNBD in the adjacent subunit. In these regions, more contact pairs are formed in the nanodisc structure (Fig. S17).

These changes cannot be attributed to overall length compaction, a phenomenon often linked with severe orientation bias in cryo-EM studies, as the overall length of HCN1 in the nanodisc is nearly identical to that in detergent. Moreover, some structural changes in the nanodisc sample involve movements inconsistent with longitudinal compaction. Another global difference observed is the slight expansion of the transmembrane domain (by 1.3Å in outer diameter) in the nanodisc structure compared to the detergent structure. This expansion may result from environmental differences and likely influences interaction forces, thereby altering allosteric coupling.

In summary, the observed changes in inter-subunit packing interactions between nanodisc- and detergent-reconstituted HCN1 channels indicates that the lipid membrane environment alters inter-subunit coupling between the transmembrane domain and neighboring C-terminal domain, resulting in increased ligand-binding cooperativity in these channels in the closed conformation.

## Discussion

In this study, we investigate the role of lipid membranes in agonist binding to ion channels by reconstituting HCN1 and HCN4 channels in nanodiscs containing either purified lipids or native membranes and compare them to detergent-solubilized channels. To differentiate between competing models of cooperativity(11, 12, 22, 62), we directly measured the microscopic equilibrium constants associated with each of the four cAMP binding events using single-molecule methods. Our results show that the binding affinity of the first cAMP molecule is unaffected by the environment—whether in detergent or a lipid bilayer. However, subsequent cAMP binding events exhibit increased affinity for HCN channels in nanodiscs, whereas these affinities remain unchanged for detergent-solubilized channels. This demonstrates that the lipid bilayer influences ligand-binding allostery, but not the intrinsic cAMP binding affinity.

To better mimic a native lipid environment and circumvent detergent exposure, we directly extracted HCN1 channels into novel lipid-encapsulated nanodiscs(59) (DeFrNDs), bypassing the detergent solubilization step. Consistent with our findings in bilayer reconstituted systems, cAMP binding to HCN1 channels in native membranes also displayed cooperative behavior, highlighting a central role of the lipid bilayer in the cellular environment. Although we cannot rule out that endogenous non-lipidic co-factors may modulate cooperativity in various tissues, the observed ligand binding cooperativity in HCN channels expressed in heterologous systems is primarily due to the lipid bilayer.

Comparison of our HCN1 structure in soy lipid nanodiscs with previously solved detergent structures reveals a plausible mechanism for bilayer-dependent cooperativity. While the overall structural features remain consistent, the nanodisc-reconstituted structure shows subtle but significant differences: the intersubunit interfaces are more tightly packed in nanodiscs, with residue-residue distances at the interface between the S4-S5 linker and CNBD C-linker being, on average, 0.9 Å shorter. These structural changes are accompanied by a 1.3 Å expansion in the outer diameter of HCN1, indicating that tighter intersubunit packing likely contributes to cooperative behavior. These observed changes are consistent with a recently published high-resolution (2.95 Å) structure of HCN1 reconstituted in PC:PE nanodiscs, further reinforcing this conclusion(61).

Although our finding that ligand binding is cooperative in HCN channels is superficially similar to a recent report (62), there are a few key differences. First and foremost, the previous study was conducted without isolating HCN channels from cellular membranes. Given that the cAMP regulation depends on the cellular context and various accessory subunits, one cannot rule out the possibility that these effects are due to interactions with specific lipids or accessory subunits(63-65). Second, significant binding of cAMP to unroofed membranes without HCN channels raises concerns about the specificity of observed binding events. Finally, these single-molecule measurements were performed at ligand concentrations exceeding the upper limit for TIRF microscopy, where it is challenging to resolve multiple binding events due to high background fluorescence (19-21). To circumvent this known limitation, the authors applied a smoothing algorithm to reduce the background signals but this approach has not been validated.

Our findings that the lipid bilayer increases intersubunit packing interactions and promotes cooperativity in HCN channels is consistent with the idea that solvating lipids in a bilayer as opposed to detergents reduce conformational flexibility, thereby destabilizing intermediate states and facilitating cooperative behavior. Membrane proteins solubilized in detergents have been found to be less stable compared to those reconstituted in nanodiscs(*61-63*). Similarly, single-molecule FRET measurements on β2-adrenergic receptors have shown that detergent-solubilized proteins undergo rapid transitions through a broad range of states in the presence of agonist, whereas receptors reconstituted in nanodiscs undergo slower transitions between active and inactive states(66).

The ability of lipid bilayers to promote cooperative binding without altering intrinsic affinity may have important physiological implications. Positive cooperativity acts as a gain factor in signaling pathways, amplifying small changes in signal while limiting the dynamic range. For HCN channels, enhancing cooperativity without changing intrinsic cAMP affinity enables lipid membranes to modulate signal amplification in pacemaker cells while preserving responsiveness to physiologically relevant cAMP concentrations. Future studies may reveal whether changes in lipid composition due to aging, metabolic shifts, or disease could modify or disrupt the integration of voltage- and cAMP-dependent signaling pathways.

## Materials and Methods

### Constructs used to express proteins used in single molecule imaging

The human ortholog of HCN4 (full length) was synthesized in the pTwist vector (Twist Bioscience) and was subcloned into the pEG BacMam vector71. From the full-length hHCN4 construct, amino acids 782-1062 (total 283 amino acids) were deleted for biochemical stability, N-terminus was tagged with eGFP followed by a 3C protease cleavage site. A twin-strep tag was introduced in the N-terminus for affinity purification to generate GFP-twinstrep-hHCN4Δ282C, this construct is HCN4SM but is referred as HCN4 from here on in the manuscript. The construct was verified by Sanger sequencing. HCN1SM was the same construct used in a previous study referred as HCN1 in this manuscript 6. Circular nanodiscs were made from spNW11 and spNW30 constructs in pET28 (generously gifted by Huan Bao laboratory), and purification of spNW11 and spNW30 was done as described in the original work(55).

### Cell Culture and Baculovirus Generation

HCN1 and HCN4 constructs described above were used to generate Baculovirus. Both constructs in the pEG vector were transformed into chemically competent DH10Bac E.coli cells (Invitrogen) as per the manufacturer’s instructions to generate bacmid DNA. One microgram of the resultant bacmid was then transfected into 1×106 cells sf9 cells (Thermo Fisher Scientific) using Cellfectin-II (Invitrogen) reagent, incubated for 72-96 hours to generate the P1 virus. The supernatant containing the P1 virus was collected 96 hours post-transfection, sterile filtered, and stored at 4C. The resultant P1 virus was further amplified by infecting sf9 cells at a dilution of 1:1000 to generate the P2 virus. The P2 virus was collected 96 hours post-transduction, sterile filtered, supplemented with 2.5% FBS (Sigma), and stored at 4°C until further use. This P2 Baculovirus was used for expressing the HCN1-SM and HCN4-SM in free-style HEK suspension cells (Thermo Fisher Scientific). An 800ml suspension of free-style HEK cells was grown to a density of 2.5×106 cells/ml for viral transduction. The cells were transduced with 4% respective P2 baculovirus. Ten to 12 hours post-transduction, 10mM sodium butyrate (Sigma Aldrich) was added to boost the channel expression71, and the flask was transferred to 30°C for another 48 hours. The culture was harvested 60 hours post-transduction by centrifugation at 3000 rpm for 20 minutes, the resultant cell pellet was flash-frozen in liquid nitrogen and stored at -80°C until further use.

### Purification and reconstitution of HCN channels into nanodiscs

Protein purification for the HCN1 detergent sample was done as reported previously6. For the nanodisc reconstitution, the same protocol was adopted with few changes to facilitate reconstitution as described here. Frozen cell pellet corresponding to 200ml cell culture of either HCN1 or HCN4 was thawed on ice and lysed with 50mL hypotonic lysis buffer (20mM KCl, 10mM Tris, protease inhibitor cocktail (Sigma Aldrich, P8340), pH 8.0) and sonicated at 4°C 5 times for 10 s each. The lysate was spun at 50,000 x g for 45 minutes to pellet the membrane fraction. The resultant membrane fraction was homogenized on ice with 20ml 2X solubilization buffer (600ml KCl, 80mM Tris, 4mM DTT, 20% glycerol, protease inhibitor cocktail (P8340, Sigma Aldrich)). After homogenization, 20ml of 2X detergent solution– 20mM Lauryl Maltose Neopentyl Glycol (LMNG, Anatrace) and 8mM Cholesteryl Hemi Succinate (CHS, Anatrace) was added to solubilize the membrane for 1.5 hours at 4°C. Later, the solubilized membranes were spun down at 50,000 x g for 45 minutes and the supernatant was incubated with pre-equilibrated Strep-Tactin Sepharose (IBA life sciences, 2-1201-025) for 2.5 hours. Then, the resin was packed onto a column and washed with 5 column volumes of wash buffer - 300 mM KCl, 20 mM Tris, 2 mM DTT, 5% glycerol, 0.5mM LMNG, 0.1mM CHS, (either 164.5 uM Soy Polar Lipids/ 197.5uM Brain polar lipids or 182uM, 3:1 POPC/POPE), pH=8.0). All the lipids were purchased from Avanti Polar Lipids as 25mg/ml solution in chloroform. The protein was eluted in 5mL elution buffer (wash buffer + 10 mM d-desthiobiotin (Sigma)) as 1ml aliquots. The eluted protein concentration was measured using a Denovix spectrophotometer and then the protein sample was concentrated to 4mg/mL using a concentrator (cut-off: 100 kDa, Amicon Ultra, Millipore).

### Reconstitution

The day before reconstitution, lipids in chloroform (Avanti Polar Lipids) were dried under argon and then desiccated overnight under a vacuum at 4°C. On the day of reconstitution, lipids were dissolved in reconstitution buffer (500mM NaCl, 50mM HEPES, 33mM CHAPS, pH 7.4) to a concentration of 10mg/ml and sonicated for 30 minutes immediately before use. Reconstitution was done by mixing purified protein, Membrane scaffolding protein (MSP), and appropriate lipid species at corresponding molar ratios (Table 1). The mixture was incubated for 30 minutes before adding 15% (v/v) Bio-beads SM2 resin (Bio-Rad) to remove the detergent. The reconstitution mix along with Biobeads was incubated overnight at 4°C on a nutator.

The next day, bio-beads were removed by brief centrifugation for 5 minutes at 2000 x g and the clear reconstitution solution was collected, concentrated to 300 µl total volume, and further purified in size exclusion chromatography (SEC) using a Superose 6 Increase 10/300 GL column (Cytiva Life Sciences, 29091596) in an SEC buffer (200 mM KCl, 20 mM Tris, 2mM DTT, pH=8.0). Representative SEC profiles and Coomassie blue-stained SDS-PAGE gels are depicted in figure S1. For the HCN4 detergent sample, all the steps until the elution were the same as described above, but the final SEC buffer contained 300 mM KCl, 20 mM Tris, 2 mM DTT, 5% glycerol, 0.1mM GDN (Anatrace). Only the peak fractions from SEC (Fig. S1) were collected and used for single molecule experiments.

### Native Nanodisc preparation

Free-style HEK suspension cells expressing HCN1 were lysed using a Branson cell disruptor. Crude membranes were isolated from cell lysate by ultracentrifugation at 184, 000 × g for 1 hour and resuspended in reconstitution buffer (50 mM Tris-HCl (pH 8), 300 mM NaCl, 10% glycerol, 2 mM DTT). Native nanodiscs were formed by incubation of crude membranes (∼10 mg/ml) with the DeFrMSP peptide Hex18A at 1mg/ml (4 °C, O/N). HCN1 native nanodiscs were isolated using Streptactin affinity resin (IBA Life Sciences) and eluted in reconstitution buffer plus 5 mM desthiobiotin. Samples were further purified on a Superdex S200 3.2/300 column in reconstitution buffer and used for single molecule experiments.

### CryoEM protein preparation

Flag-HCN1-TGP-10His channels, the human HCN1 1-636 residues with N-terminal flag tag and a C-terminal fusion of the thermostable green protein (TGP) and 10X His tag, were expressed in yeast (P. pastoris strain SMD1163H, Invitrogen, #C17500). The yeast cultures were grown for 48 hours post-induction, and the resultant cell pellet expressing HCN1 was disrupted by milling (Retsch MM400). The ground powder was resuspended in a buffer containing 50mMTris, 300mM KCl, 10%Glycerol, 1mM EDTA with protease inhibitor cocktail (Sigma) and homogenized. The homogenate was centrifuged at 100,000 x g for 40 minutes at 4°C. The supernatant was discarded, and the membrane pellet was resuspended in 2X solubilization buffer (80 mM Tris, 600mM KCl, 8mM DTT, 2x protease inhibitor cocktail, 10% glycerol, pH8.0) and flash frozen in liquid nitrogen and stored at -80°C until further use. On the day of preparation, yeast cell membranes equivalent to 10g of pellet was thawed on ice for 30 minutes on ice. Membrane fraction was then solubilized by adding an equal volume of 10 mM LMNG and 2mM CHS solution into membrane solution and incubated for 1 hour 30 minutes at 4°C on a nutator. The solubilized membrane was centrifuged at 40,000 x g for 45 minutes. The supernatant was added to Ni-NTA resin (QIAGEN) pre-equilibrated with 200mM KCl, 20mM Tris, 2mM TCEP, 0.05% LMNG-CHS, 160µM soy polar lipids, and incubated for 90 minutes at 4°C on a nutator. After batch binding, the resin with the supernatant was loaded onto a column and allowed to pack under gravity. The resin was washed with wash buffer (200mM KCl, 20mM Tris, 2mM TCEP, 0.05% LMNG-CHS, 160µm soy polar lipids (3X column volumes). Protein was eluted with elution buffer (SEC buffer with 250mM imidazole), and the protein concentration was measured using a Denovix spectrophotometer. The affinity-purified protein was concentrated using a 100 KDa-cutoff concentrator and further purified in size exclusion chromatography (SEC) using a Superose 6 Increase 10/300 GL column (Cytiva Life Sciences, 29091596) in a SEC buffer (200 mM KCl, 20 mM Tris, 0.0025% LMNG, 62.5 µM CHS, 7.5 µM soy lipid mix, pH 8.0). The peak fraction from SEC was collected and then concentrated using a 100KDa concentrator. Protein was reconstituted with MSP2N2, soy polar lipid at a 1:2.5:200 molar ratio. The mixture was incubated for 30 minutes before adding 15% (v/v) Bio-beads SM2 resin (Bio-Rad) to remove the detergent. The reconstitution mix along with Biobeads was incubated overnight at 4°C on a nutator. The next day, bio-beads were removed by brief centrifugation for 5 minutes at 2000xg and the clear reconstitution solution was collected. This was then incubated with anti-Flag resin (Genscript Anti-DYKDDDDK G1 Affinity Resin, Cat#L00432) preequilibrated with 200 mM KCl, 20 mM Tris, 2mM DTT, pH=8.0 for a second round of purification to remove any empty nanodiscs. The flag resin-bound nanodiscs were eluted with 200 mM KCl, 20 mM Tris, 2 mM DTT, 250ug/ml of Flag peptide (Sigma Cat #F3290) pH=8.0. The eluate was concentrated to 300µl and further purified in size exclusion chromatography (SEC) using a Superose 6 Increase 10/300 GL column (Cytiva Life Sciences, 29091596) in an SEC buffer (200 mM KCl, 20 mM Tris, 2mM DTT, pH=8.0).

### CryoEM Data Acquisition and Analysis

The purified nanodisc with HCN1 at 2.0mg/ml concentration was frozen in Quantifoil R1.2/1.3 copper grids with 300 mesh with Thermo Fisher Vitrobot Mk IV in the Center for Cellular Imaging at Washington University in St. Louis (WUCCI). The cryoEM imaging was performed in WUCCI with the ThermoFisher Scientific Glacios 200kV Cryo-TEM. The imaging parameters are listed in the Table 2. Un-tilted imaging results without surfactant showed that the particles are mainly in top/bottom orientation, which prevents getting correct cryoEM map without compression. Adding 1 mM fluorinated fos-choline to the sample resulted in aggregation. Non-tilted imaging with 0.2mM fluorinated fos-choline did not have a major improvement in particle orientation. The cryoEM imaging with 30° tilt for 24 hours (without surfactant) resulted in 768 micrographs. Single particle analysis was performed with cryosparc v4.4.1 mainly from tilted imaging results combined with some particles from non-tilted imaging. The detailed flowchart for analysis is shown in figure S14. This resulted in a final map with 3.77Å resolution.

**Table 2.** Imaging parameters for CryoEM data acquisition.

|  | Dataset 1 | Dataset2 | Dataset3 |
| --- | --- | --- | --- |
| Surfactant | no | no | 0.2mM FI-fos-choline |
| Tilting (°) | 30 | 30 | 0 |
| Accelerating Voltage (kV) | 200 | 200 | 200 |
| Pixel Size (Å) | 1.184 | 1.184 | 1.184 |
| Spherical Aberration (mm) | 2.7 | 2.7 | 2.7 |
| Total Dose (e/Å<sup>2</sup>) | 47.9 | 43.13 | 50.63 |
| Fractions | 40 | 40 | 45 |
| Number of micrographs | 279 | 489 | 6993 |

The model building started with fitting alphafold model AF-A0A669KB45-F1-model_v4 to the density in Chimerax 1.6, and then manually adjusted in coot v0.9.8.93. After validation of the monomer, the tetramer was built with Chimera 1.17.3 with symmetry expansion to match the C4 symmetry in the map. The tetramer was further real-space refined with Phenix, and the resulting model was further adjusted to reduce clashes. The final model was validated in the PDB validation server (https://validate-rcsb-1.wwpdb.org/).

Morph from the hHCN1-detergent model (5u6o) to the hHCN1-nanodisc model was performed in Chimerax 1.6 by fitting the detergent map (emd_8511) to the nanodisc map first and then fitting models to their own maps to superimpose two models before morphing.

Residue-residue distance (Cα-Cα distances) was generated by chimera RR distance map with two concatenated subunits, making it possible to get intersubunit 2D contact map. The RR distance was exported to a CSV file, which was opened in Excel, and processed with IFS function to generate a contact matrix with 10 Å cutoff. The final 2D contact matrix in the subunit interface was copied and pasted to the matrix in OriginLab and plotted as heatmap with distance information within 10Å. Sub-regions in the 2D intersubunit contact map were further selected and plotted to show more detail residue-residue contacts.

2D contact maps with the same cutoff were also generated in Discovery Studio Visualizer 2024. Selecting a sub-region in the 2D contact map resulted in selection of corresponding residues in the structural model. Thus, we use this feature to plot contacting residues identified in the 2D contact map to the 3D structure. The final figure was arranged in Affinity Designer 2.

### Zero mode waveguides Fabrication

ZMW fabrication was done at The Center for Nanophase Materials Sciences, Oak Ridge National Laboratory. Briefly, Cover glasses (Fisher Scientific cat. no.12-548C, 130-170 um thickness) were cleaned with 2% Hellmanex (Hellma), HPLC grade Acetone (Sigma), HPLC grade Ethanol (Sigma), and 3 M KOH by sonicating for 15 minutes in each solution, intervened by a water wash. Cleaned cover glasses were plasma cleaned (Gatan Solarus 950 plasma cleaner) for 10 minutes. Cover glasses were coated with aluminum using either sputter coating at a 4 nm/min rate using a DC Magnetron sputtering or thermally evaporated aluminum at a rate of 2 Å/second using a JEOL dual source E-beam evaporator for a total thickness of 100 nm. In DC sputtering, an argon plasma is used to initiate deposition. Excited argon ions strike the target material, causing atoms to be ejected from the target. These atoms then deposit onto the substrate, which is placed below the target source. On the other hand, the E-beam evaporator utilizes an electron beam to heat metals until they reach a vapor phase. Substrates are positioned above the electron beam guns, and when the shutter is opened, the vaporized metal is deposited onto the substrates. The deposition thickness was checked with a surface profilometer and confirmed to be 100 ± 5 nm. Aluminum-coated cover glasses were then spin-coated with positive-tone electron beam resist ZEP 520A (45 sec at 3000 rpm). The cover glasses were baked at 180 °C for 2 mins. After that, ZMWs featuring 120, 150, and 180 nm dots were patterned on the surface using 100 kV accelerating voltage and 2 nA beam current using the JEOL JBX-8100FS electron beam lithography system. Following exposure, the ZMWs were developed using a 30-sec treatment of xylenes, followed by a wash with 2-propanol, and dried with nitrogen gas. Developed cover glasses were mounted on a silicon wafer using a thin layer of Krytox oil applied on the back of the cover glasses, and 100 nm aluminum was dry etched for 1 min using OXFORD Plasmalab System 100 reactive ion etcher with a mixture of 30 standard cubic centimeters (sccm) chlorine (Cl2) and 10 sscm boron trichloride (BCl3) gasses at 50 °C. Finally, the samples were plasma cleaned in PVA TePla IoN Wave 10 using O2-Ar plasma to remove any excess photoresist. To ensure the quality, ZMWs from different batches were chosen at random and imaged under Zeiss Merlin FE-Scanning Electron Microscopy (SEM). We note that the hole sizes were within the margin of error ±10 nm.

### Sample and chamber preparation for single-molecule imaging

Cover glasses and ZMWs used for single-molecule imaging were cleaned and passivated as described previously6. Briefly, ZMWs were plasma cleaned, treated with 2% polyvinyl phosphonic acid (Polysciences), washed with water, and dried under argon gas. Later the ZMW was passivated with mPEG-silane (Laysan Bio) and doped with biotin-PEG silane (Laysan Bio) overnight. A small 50 µl silicone-gasketed microfluidic chamber (Grace Bio labs) was attached to the ZMW for single-molecule imaging. Single-molecule pulldown was done to immobilize the protein. Briefly, the chamber was sequentially incubated with 10% BSA (Sigma Aldrich) for 30 minutes (additional layer of passivation), 1 µM streptavidin (Prospec) for 10 minutes, 10 nM biotinylated GFP nanobody (Chromtek) for 10 minutes, ∼225-250 pM of GFP-tagged-HCN1 and HCN4 for 10 minutes. After 10 minutes, the chambers were washed to remove excess freely diffusing protein and empty nanodiscs. Each of the steps before was also intervened by a wash step with 1X Tris-buffered saline to remove excess freely diffusing molecules. All the single-molecule experiments were done in an imaging buffer (1X Tris-buffered saline supplemented with 1% BSA, 2 mM Dithiothreitol (DTT, Sigma-Aldrich) and oxygen scavengers including 2mM Trolox (Sigma-Aldrich) and 2.5mM protocatechuic acid (Sigma-Aldrich) and 250nM protocatechuate 3, 4-dioxygenase from Pseudomonas sp. (Sigma-Aldrich) were added just before imaging. If the sample being imaged was a protein in the detergent micelle, then the buffer contained 100uM Glycodiosgenin (GDN). The chamber was then added with various concentrations of 8-(2-[DY-547}-aminoethylthio) adenosine-3’, 5’-cyclic monophosphate (fcAMP; Biolog). The solutions in the chamber were replenished every 20 minutes to avoid evaporation.

### Single-molecule imaging

Imaging was done on an inverted microscope (Mad City Labs) on a high NA 60X oil immersion objective lens (Olympus) under a micromirror TIRF excitation field generated either with a 488nm or 561nm lasers (OBIS, coherent) to excite the GFP and fcAMP respectively. The subsequent fluorescence emission was recorded on a 512×512 EMCCD camera (Andor iXON Ultra) at a frame rate of 10Hz. Data collection was accomplished using Micromanager 2.0. The use of micromirror TIRF eliminated most of the stray excitation light. The emission was passed through a dichroic filter (T565lpxr Chroma) and then a band-pass filter (Chroma ET550/25 nm) for GFP and a band-pass filter (Semrock bright line 593/40nm) for fcAMP were in the corresponding emission pathways. The micromirror TIRF is set up for a 160X magnification and 234×234um field of view, allowing us to view the ∼720 ZMWs at once. Diameter of the input laser beam was optimized to 5mm which gives a 100uM illumination spot on the objective and the laser power measured using an optical power meter (Newport, 1918-R) at the objective was 9.6mW and 7mW for 488nm and 561nm lasers respectively. The micromirror TIRF setup was equipped with a TIRF-lock that continuously monitors the position of the TIRF exit beam and adjusts the Nano-positioner for any drift in the Z-axis48,72, MWs were labeled sparsely to reduce the probability of having more than one per ZMW. ZMWs that are labeled were identified by exciting the GFP molecules, these GFP molecules were photo-bleached. Later, fcAMP molecules were excited with the 561nm laser to measure the binding activity. The binding data were recorded from each molecule for a minimum of 240 seconds with 100 msec exposure time. Single-molecule data that was collected was analyzed exactly as described in the previous work6 with the exception that instead of DISC6,26 AutoDISC49 was used to idealize the traces for the unsupervised selection of objective criteria during the data analysis. Briefly, single-molecule traces were extracted and analyzed using the DISC software for the colocalization and image projection to obtain time-dependent fluorescence intensity changes, the traces were then idealized in autoDISC. All the idealized traces were visually inspected to eliminate molecules with more than 4 bound states and traces with low signal-to-noise where state assignment was difficult. The data analysis was repeated to remove any single frame events arising from dye diffusion, blinking, noise, etc. which may have been identified as bound states. From the Idealized Fluorescence-time traces, we extracted the time spent by fcAMP in each ligation state (state occupancy) at every. These state occupancies were normalized to the Expected value obtained from a Binomial distribution assuming independent and identical binding sites, each with a probability of ligand binding P, the probability of x binding sites being occupied simultaneously is given by the equation:

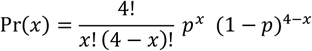

The ratio of Observed over Expected occupancies is 1 when the ligand binding is completely independent. All the data analysis was done using custom MATLAB (Mathworks) scripts. HMM, analysis of all the data sets was performed using QuB as described before(23). Briefly, for each set of conditions, models were globally optimized using maximum idealized point likelihood rate estimation (MIP) across all fcAMP concentrations. The goodness of fit was measured by the Bayesian Information Criterion (BIC). BIC = k X ln (n) - 2 x LL.

The microscopic rate constants were extracted from the following model, in which each of the microscopic equilibrium constants was allowed to float freely. The empty squares represent unoccupied cAMP binding sites, gray circles represent fcAMP, and filled squares represent occupied cAMP binding sites.

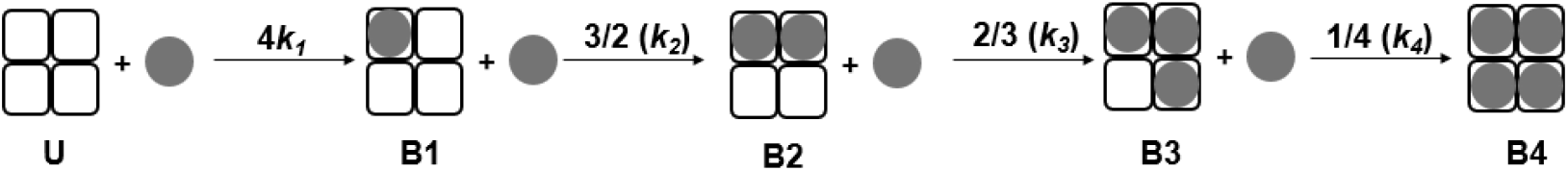

The extrapolated binding curve was generated using the microscopic rate constants extracted from QuB and plugging into the Adair’s equation

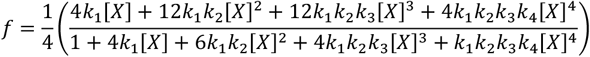

Where ‘f ‘ is the bound probability; k1, k2, k3, and k4 are association rate constants for 1st, 2nd, 3rd and 4th bound states respectively; and [X] is the fcAMP concentration. Note that, a bound probability of 1.0 corresponds to fcAMP occupying all four binding sites.

## Acknowledgments

This research was funded by grants from the National Institutes of Health to B.C. (NS116850) and H.B. (DP2GM140920) and by the National Science Foundation to R.H.G. (CHE-1856518). The fabrication of the ZMW waveguides was conducted as part of a user project at the Center for Nanophase Materials Sciences (CNMS), a US Department of Energy, Office of Science User Facility at Oak Ridge National Laboratory. The authors would like to acknowledge guidance and training in the fabrication process by Bernadeta Srijanto, Dale Hensley, Daryl Briggs, Kevin Lester, Leslie Wilson, and Scott Retterer. Cryo-EM data were collected at the Washington University Center for Cellular Imaging (WUCCI), supported by the Washington University School of Medicine, The Children’s Discovery Institute of Washington University, and St. Louis Children’s Hospital (CDI-CORE-2015-505 and CDI-CORE-2019-813) and the Foundation for Barnes-Jewish Hospital (3770 and 4642). We thank Drs. B. Summers and K. Basore at WUCCI for grid freezing and help with cryo-EM data collection. We thank Drs. W. Cheng and A.S. Evers for discussion and comments.

## References

1. C. Wahl-Schott, M. Biel, HCN channels: structure, cellular regulation and physiological function. Cellular and molecular life sciences 66, 470–494 (2009).

2. R. B. Robinson, S. A. Siegelbaum, Hyperpolarization-activated cation currents: from molecules to physiological function. Annual review of physiology 65, 453–480 (2003).

3. A. Ludwig, X. Zong, M. Jeglitsch, F. Hofmann, M. Biel, A family of hyperpolarization-activated mammalian cation channels. nature 393, 587–591 (1998).

4. R. Gauss, R. Seifert, U. B. Kaupp, Molecular identification of a hyperpolarization-activated channel in sea urchin sperm. Nature 393, 583–587 (1998).

5. B. Santoro et al., Identification of a gene encoding a hyperpolarization-activated pacemaker channel of brain. Cell 93, 717–729 (1998).

6. D. DiFrancesco, Pacemaker mechanisms in cardiac tissue. Annual review of physiology 55, 455–472 (1993).

7. R. Gauss, R. Seifert, Pacemaker oscillations in heart and brain: a key role for hyperpolarization-activated cation channels. Chronobiology international 17, 453–469 (2000).

8. D. DiFrancesco, P. Tortora, Direct activation of cardiac pacemaker channels by intracellular cyclic AMP. Nature 351, 145–147 (1991).

9. V. Nache et al., Activation of olfactory-type cyclic nucleotide-gated channels is highly cooperative. The Journal of physiology 569, 91–102 (2005).

10. J. W. Karpen, A. L. Zimmerman, L. Stryer, D. A. Baylor, Gating kinetics of the cyclic-GMP-activated channel of retinal rods: flash photolysis and voltage-jump studies. Proceedings of the National Academy of Sciences 85, 1287–1291 (1988).

11. J. Kusch et al., Interdependence of receptor activation and ligand binding in HCN2 pacemaker channels. Neuron 67, 75–85 (2010).

12. J. Kusch et al., How subunits cooperate in cAMP-induced activation of homotetrameric HCN2 channels. Nature chemical biology 8, 162–169 (2012).

13. D. T. Liu, G. R. Tibbs, P. Paoletti, S. A. Siegelbaum, Constraining ligand-binding site stoichiometry suggests that a cyclic nucleotide–gated channel is composed of two functional dimers. Neuron 21, 235–248 (1998).

14. J. Li, H. A. Lester, Single-channel kinetics of the rat olfactory cyclic nucleotide-gated channel expressed in Xenopus oocytes. Molecular pharmacology 55, 883–893 (1999).

15. J. Li, W. N. Zagotta, H. A. Lester, Cyclic nucleotide-gated channels: structural basis of ligand efficacy and allosteric modulation. Quarterly reviews of biophysics 30, 177–193 (1997).

16. M. Lolicato et al., Tetramerization dynamics of C-terminal domain underlies isoform-specific cAMP gating in hyperpolarization-activated cyclic nucleotide-gated channels. Journal of Biological Chemistry 286, 44811–44820 (2011).

17. K. E. Hines, T. R. Middendorf, R. W. Aldrich, Determination of parameter identifiability in nonlinear biophysical models: A Bayesian approach. Journal of General Physiology 143, 401–416 (2014).

18. T. R. Middendorf, M. P. Goldschen-Ohm, The surprising difficulty of “simple” equilibrium binding measurements on ligand-gated ion channels. Journal of General Physiology 154, e202213177 (2022).

19. D. S. White, M. A. Smith, B. Chanda, R. H. Goldsmith, Strategies for Overcoming the Single-Molecule Concentration Barrier. ACS Measurement Science Au 3, 239–257 (2023).

20. S. Peng, W. Wang, C. Chen, Breaking the Concentration Barrier for Single-Molecule Fluorescence Measurements. Chemistry–A European Journal 24, 1002–1009 (2018).

21. P. Holzmeister, G. P. Acuna, D. Grohmann, P. Tinnefeld, Breaking the concentration limit of optical single-molecule detection. Chemical Society Reviews 43, 1014–1028 (2014).

22. D. S. White et al., cAMP binding to closed pacemaker ion channels is non-cooperative. Nature 595, 606–610 (2021).

23. D. S. White, M. P. Goldschen-Ohm, R. H. Goldsmith, B. Chanda, Top-down machine learning approach for high-throughput single-molecule analysis. Elife 9, e53357 (2020).

24. M. J. Levene et al., Zero-mode waveguides for single-molecule analysis at high concentrations. science 299, 682–686 (2003).

25. P. Zhu, H. G. Craighead, Zero-mode waveguides for single-molecule analysis. Annual review of biophysics 41, 269–293 (2012).

26. J. Chen et al., High-throughput platform for real-time monitoring of biological processes by multicolor single-molecule fluorescence. Proceedings of the National Academy of Sciences 111, 664–669 (2014).

27. M. P. Goldschen-Ohm, D. S. White, V. A. Klenchin, B. Chanda, R. H. Goldsmith, Observing Single-Molecule Dynamics at Millimolar Concentrations. Angewandte Chemie 129, 2439–2442 (2017).

28. S. Uemura et al., Real-time tRNA transit on single translating ribosomes at codon resolution. Nature 464, 1012–1017 (2010).

29. J. Hentschel et al., Real-time detection of human telomerase DNA synthesis by multiplexed single-molecule FRET. Biophysical journal 122, 3447–3457 (2023).

30. P. Pian, A. Bucchi, R. B. Robinson, S. A. Siegelbaum, Regulation of gating and rundown of HCN hyperpolarization-activated channels by exogenous and endogenous PIP2. The Journal of general physiology 128, 593–604 (2006).

31. P. A. Schmidpeter et al., Anionic lipids unlock the gates of select ion channels in the pacemaker family. Nature structural & molecular biology 29, 1092–1100 (2022).

32. C. J. daCosta, L. Dey, J. P. Therien, J. E. Baenziger, A distinct mechanism for activating uncoupled nicotinic acetylcholine receptors. Nat Chem Biol 9, 701–707 (2013).

33. R. Rusinova, C. He, O. S. Andersen, Mechanisms underlying drug-mediated regulation of membrane protein function. Proc Natl Acad Sci U S A 118 (2021).

34. I. Levental, E. Lyman, Regulation of membrane protein structure and function by their lipid nano-environment. Nat Rev Mol Cell Biol 24, 107–122 (2023).

35. B. Reddy, N. Bavi, A. Lu, Y. Park, E. Perozo, Molecular basis of force-from-lipids gating in the mechanosensitive channel MscS. Elife 8 (2019).

36. E. Perozo, D. M. Cortes, P. Sompornpisut, A. Kloda, B. Martinac, Open channel structure of MscL and the gating mechanism of mechanosensitive channels. Nature 418, 942–948 (2002).

37. Y. Zhang et al., Visualization of the mechanosensitive ion channel MscS under membrane tension. Nature 590, 509–514 (2021).

38. H. Zheng, W. Liu, L. Y. Anderson, Q.-X. Jiang, Lipid-dependent gating of a voltage-gated potassium channel. Nature communications 2, 250 (2011).

39. C. Yuan, R. J. O’Connell, R. F. Jacob, R. P. Mason, S. N. Treistman, Regulation of the gating of BKCa channel by lipid bilayer thickness. Journal of Biological Chemistry 282, 7276–7286 (2007).

40. H. X. Zhou, T. A. Cross, Influences of membrane mimetic environments on membrane protein structures. Annu Rev Biophys 42, 361–392 (2013).

41. C. R. Sanders, K. F. Mittendorf, Tolerance to changes in membrane lipid composition as a selected trait of membrane proteins. Biochemistry 50, 7858–7867 (2011).

42. V. R. Patel et al., Single-molecule imaging with cell-derived nanovesicles reveals early binding dynamics at a cyclic nucleotide-gated ion channel. Nature Communications 12, 6459 (2021).

43. M. P. Goldschen-Ohm et al., Structure and dynamics underlying elementary ligand binding events in human pacemaking channels. Elife 5, e20797 (2016).

44. J. Larson et al., Design and construction of a multiwavelength, micromirror total internal reflectance fluorescence microscope. Nature protocols 9, 2317–2328 (2014).

45. A. Bandyopadhyay, M. P. Goldschen-Ohm, Unsupervised selection of optimal single-molecule time series idealization criterion. Biophysical Journal 120, 4472–4483 (2021).

46. G. S. Adair, A. Bock, H. Field Jr, The hemoglobin system: VI. The oxygen dissociation curve of hemoglobin. Journal of Biological Chemistry 63, 529–545 (1925).

47. C.-L. Huang, S. Feng, D. W. Hilgemann, Direct activation of inward rectifier potassium channels by PIP2 and its stabilization by Gβγ. Nature 391, 803–806 (1998).

48. N. Gamper, M. S. Shapiro, Regulation of ion transport proteins by membrane phosphoinositides. Nature Reviews Neuroscience 8, 921–934 (2007).

49. T. Rohacs, Phosphoinositide regulation of TRP channels. Mammalian Transient Receptor Potential (TRP) Cation Channels: Volume II, 1143–1176 (2014).

50. D. W. Hilgemann, S. Feng, C. Nasuhoglu, The complex and intriguing lives of PIP2 with ion channels and transporters. Science’s STKE 2001, re19–re19 (2001).

51. A.A. Rodríguez-Menchaca, S. K. Adney, L. Zhou, D. E. Logothetis, Dual regulation of voltage-sensitive ion channels by PIP2. Frontiers in pharmacology 3, 170 (2012).

52. M. Kruse, G. R. Hammond, B. Hille, Regulation of voltage-gated potassium channels by PI (4, 5) P2. Journal of general physiology 140, 189–205 (2012).

53. V. Dalal et al., Lipid nanodisc scaffold and size alter the structure of a pentameric ligand-gated ion channel. Nature communications 15, 25 (2024).

54. S. Ganapathy et al., Membrane matters: The impact of a nanodisc-bilayer or a detergent microenvironment on the properties of two eubacterial rhodopsins. Biochimica et Biophysica Acta (BBA)-Biomembranes 1862, 183113 (2020).

55. S. Zhang et al., One-step construction of circularized nanodiscs using SpyCatcher-SpyTag. Nature communications 12, 5451 (2021).

56. I. Schachter, C. Allolio, G. Khelashvili, D. Harries, Confinement in nanodiscs anisotropically modifies lipid bilayer elastic properties. The Journal of Physical Chemistry B 124, 7166–7175 (2020).

57. M. Daniilidis, M. J. Brandl, F. Hagn, The advanced properties of circularized MSP nanodiscs facilitate high-resolution NMR studies of membrane proteins. Journal of Molecular Biology 434, 167861 (2022).

58. A. Moroni et al., Hyperpolarization-activated cyclic nucleotide-gated channel 1 is a molecular determinant of the cardiac pacemaker current I f. Journal of Biological Chemistry 276, 29233–29241 (2001).

59. Q. Ren et al., DeFrND: detergent-free reconstitution into native nanodiscs with designer membrane scaffold peptides.. bioRxiv 10.1101/2024.11.07.622281 (2024).

60. C.-H. Lee, R. MacKinnon, Structures of the human HCN1 hyperpolarization-activated channel. Cell 168, 111-120. e111 (2017).

61. E. D. Kim et al., Propofol rescues voltage-dependent gating of HCN1 channel epilepsy mutants. Nature 632, 451–459 (2024).

62. S. Kuschke et al., cAMP binding to closed pacemaker ion channels is cooperative. Proceedings of the National Academy of Sciences 121, e2315132121 (2024).

63. C. H. Peters et al., Isoform-specific regulation of HCN4 channels by a family of endoplasmic reticulum proteins. Proceedings of the National Academy of Sciences 117, 18079–18090 (2020).

64. C. H. Peters, R. K. Singh, J. R. Bankston, C. Proenza, Regulation of HCN channels by protein interactions. Frontiers in Physiology 13, 928507 (2022).

65. Z. Liao et al., Cellular context and multiple channel domains determine cAMP sensitivity of HCN4 channels: ligand-independent relief of autoinhibition in HCN4. Journal of General Physiology 140, 557–566 (2012).

66. R. Lamichhane et al., Single-molecule view of basal activity and activation mechanisms of the G protein-coupled receptor β2AR. Proceedings of the National Academy of Sciences 112, 14254–14259 (2015).

