## Supporting information for "Lipid bilayers determine allostery but not intrinsic affinity of cAMP binding to pacemaker channels"

**This PDF file includes:**

Figs. S1 to S17
Tables S1 to S35

**Other supporting materials for this manuscript include the following:**

Movies S1 to S2

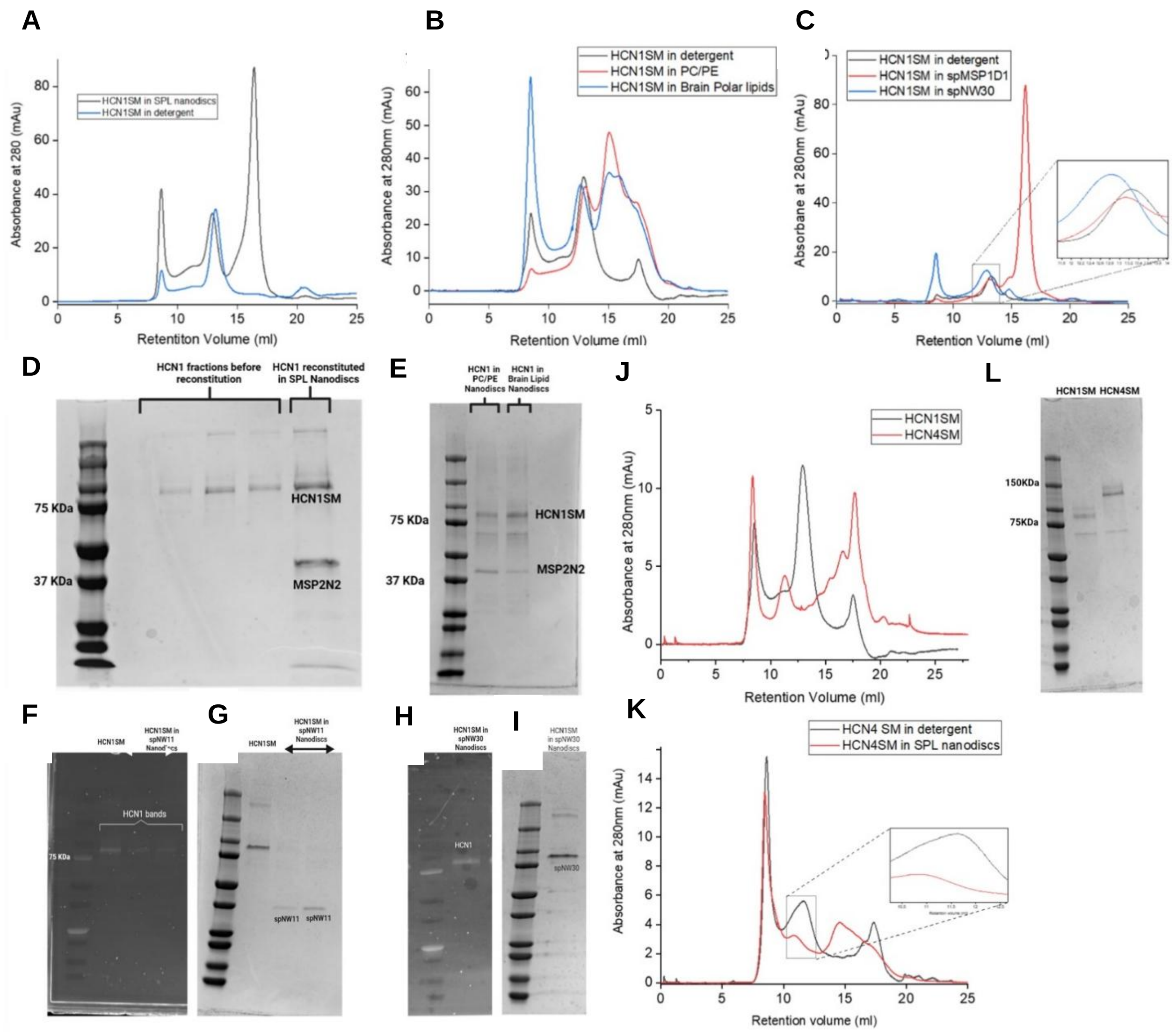

**Fig. S1. Purification and reconstitution of HCN1 and HCN4 in various lipid nanodiscs. (A)** Comparison of size exclusion chromatography profile of HCN1 purified in detergent (blue) and HCN1 purified in detergent and reconstituted into soy polar lipid nanodiscs with MSP2N2 scaffolding protein (black). **(B)** Comparison of size exclusion chromatography profile of HCN1 purified in detergent (black) and HCN1 purified in detergent and reconstituted into either Brain polar lipid nanodiscs with MSP2N2 scaffolding protein (blue) or POPC-POPE lipid nanodiscs with MSP2N2 as scaffolding protein (red). **(C)** Comparison of size exclusion chromatography profile of HCN1 purified in detergent (black) and HCN1 purified in detergent and reconstituted into brain polar lipid nanodiscs either with spMSP1D1 (11 nm nanodiscs) (red) or spNW30 (30 nm nanodiscs) (blue) scaffolding protein. **(D)** Coomassie blue stained SDS-PAGE gel showing various SEC fractions of HCN1 at ~75 kDa before reconstitution (lanes-2, 3, 4) and both HCN1 and MSP bands after reconstitution (lane-5). **(E)** Coomassie blue stained SDS-PAGE gel of HCN1 reconstituted into POPC-POPE nanodiscs (lane-2) or brain polar lipid nanodiscs (lane-3) showing both HCN1 and MSP bands. **(F)** Fluorescence image of SDS-PAGE gel showing fluorescence from GFP-HCN1 **(G)** Corresponding Coomassie blue stained SDS-PAGE gel showing spMSP1D1 band. **(H)** Fluorescence image of SDS-PAGE gel showing fluorescence from GFP-HCN1 **(I)** Corresponding Coomassie blue stained SDS-PAGE gel showing spNW30 band. **(J)** Comparison of size exclusion chromatography profile of HCN1 purified in detergent (black) and HCN4 purified in detergent (red). **(K)** Comparison of size exclusion chromatography profile of HCN4 purified in detergent (black) and HCN4 purified in detergent and reconstituted into soy polar lipid nanodiscs (red). **(L)** Coomassie blue stained SDS-PAGE gel showing HCN1 band at ~75 kDa and HCN4 band at ~100 kDa.

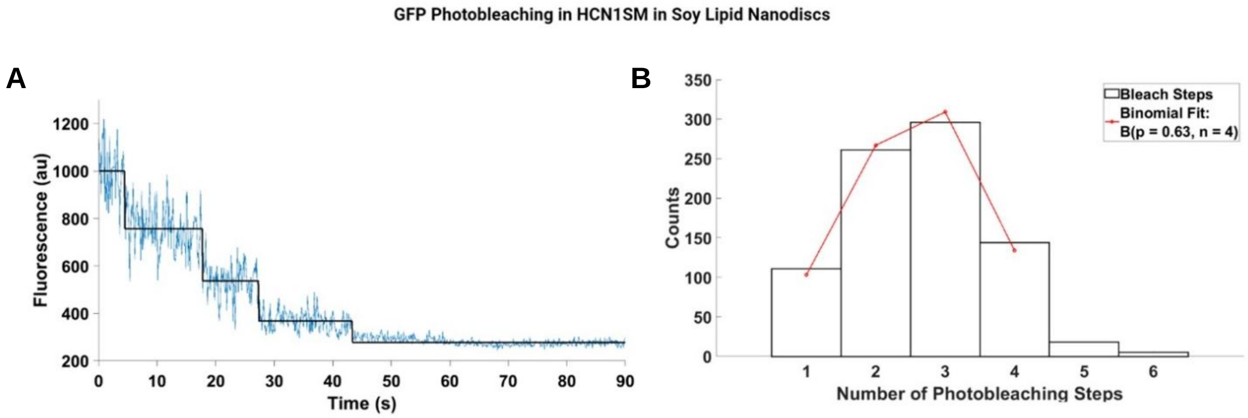

**Fig.S2. Photobleaching steps of HCN1 in soy polar lipid nanodiscs. (A)** Example photobleaching trajectory of eGFP-tagged HCN1 nanodisc tetramers on a cover glass with four identified photobleaching steps overlaid with the idealized fit (black). **(B)** Distribution of observed number of photobleaching steps of HCN1 nanodiscs overlaid with maximum likelihood estimations of a zero-truncated binomial distribution ($N$ = 835, $B$ ($n$ = 4, $p$ = 0.63).

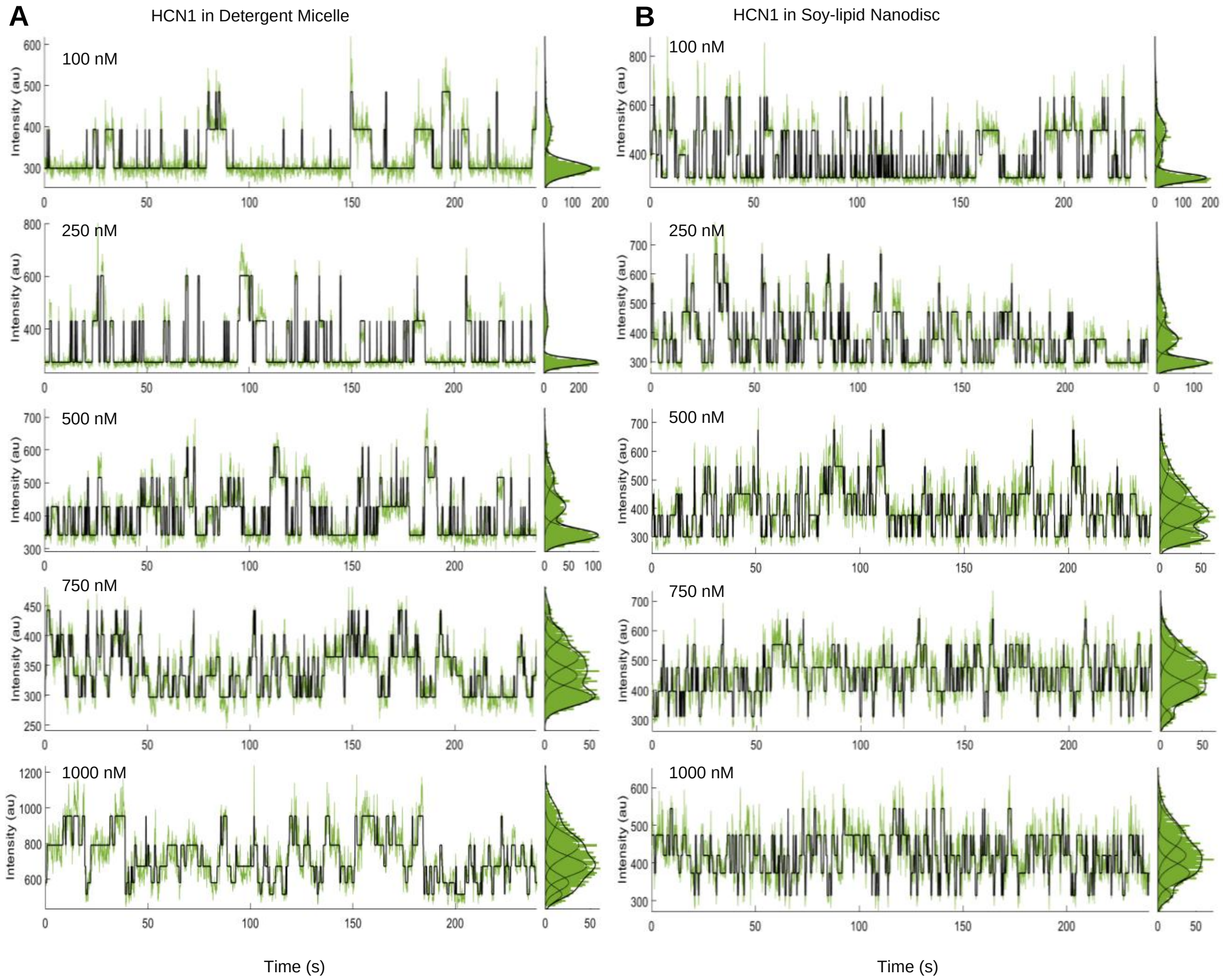

**Fig. S3. Representative fluorescence-time traces for HCN1 in detergent micelle and HCN1 reconstituted in soy polar nanodiscs. (A)** Representative fluorescence-time traces for fcAMP binding to HCN1 in detergent micelle overlaid with the idealized fit (black). Traces are shown for 100nM, 250 nM, 500 nM, 750 nM and 1000 nM fcAMP concentrations. **(B)** Representative fluorescence-time traces for fcAMP binding to HCN1 reconstituted into soy polar lipid nanodiscs overlaid with the idealized fit (black). Traces are shown for 100 nM, 250 nM, 500 nM, 750 nM and 1000 nM fcAMP concentrations.

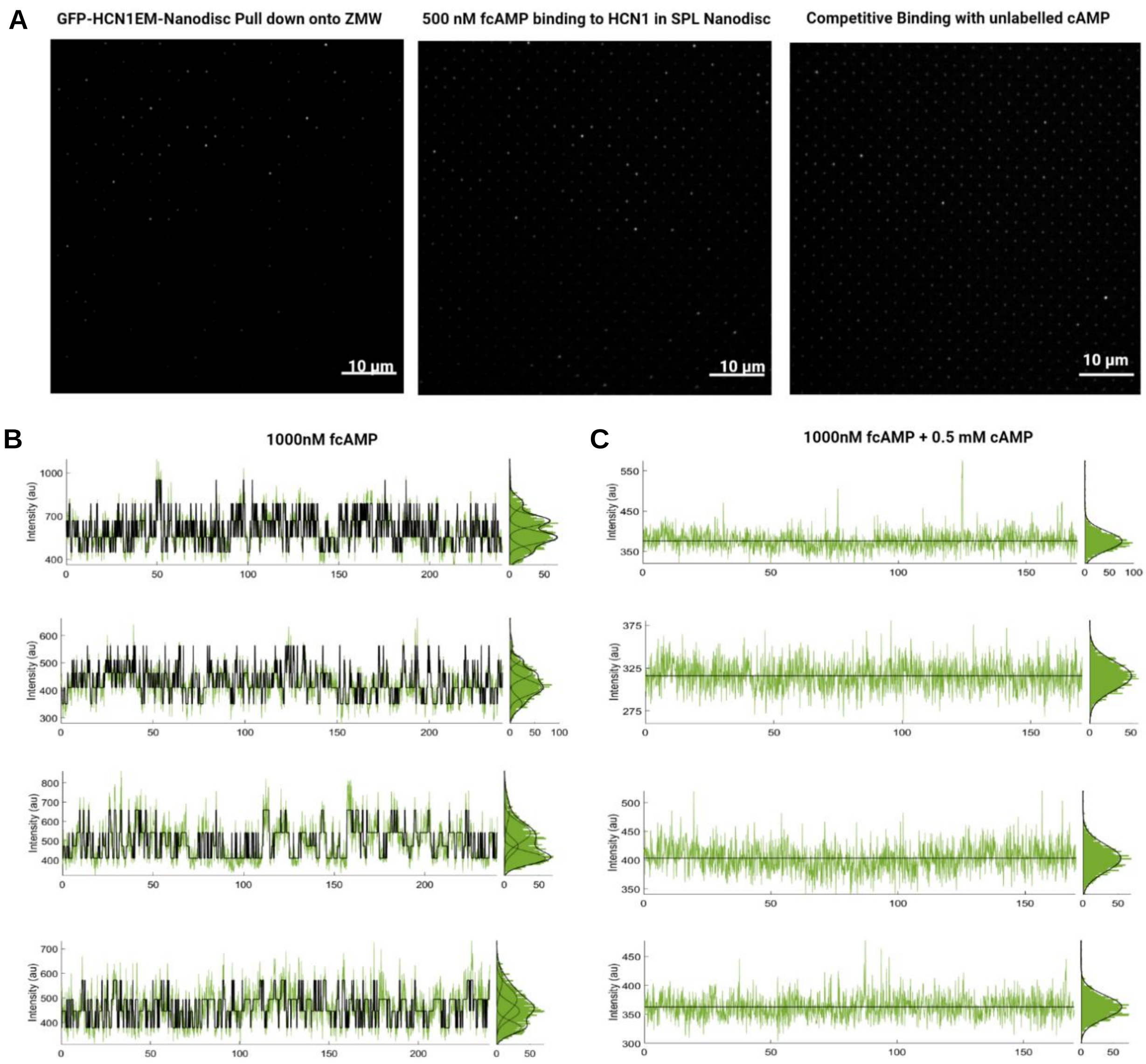

**Fig. S4. Specific binding of HCN1 to surface-modified ZMWs and fcAMP to HCN1 in ZMWs. (A)** TIRF image of ZMW array showing GFP fluorescence (488 nm excitation) from HCN1 nanodisc (left); TIRF image of ZMW array showing 500 nM fcAMP binding (561nm excitation) to HCN1 nanodiscs (middle) and TIRF image of ZMW array showing loss of fcAMP binding (561nm excitation) to HCN1 nanodiscs after addition of unlabeled cAMP (right). All the images were background subtracted for visualization. Brightness and contrast were adjusted for clarity. **(B)** Representative and randomly selected fluorescence trajectories with idealized fit (black) showing 500 nM fcAMP binding to HCN1SM nanodiscs **c**, Fluorescence trajectories from corresponding molecules showing loss of binding activity upon addition of excess unlabeled cAMP.

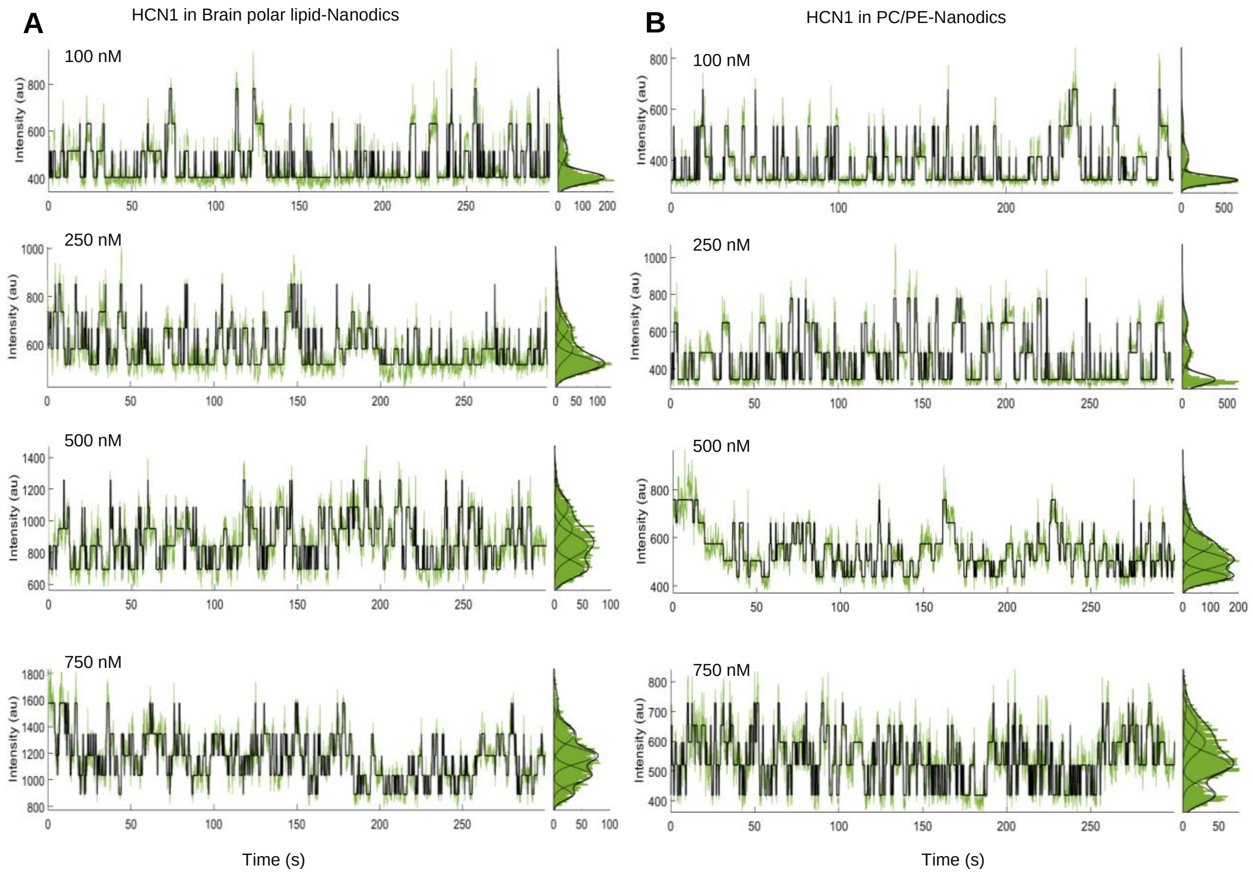

**Fig. S5. Representative fluorescence-time traces for HCN1 in brain polar lipid nanodiscs and HCN1 reconstituted in POPC-POPE nanodiscs. (A)** Representative fluorescence-time traces for fcAMP binding to HCN1 in brain polar lipid nanodiscs overlaid with the idealized fit (black). Traces are shown for 100 nM, 250 nM, 500 nM, 750 nM fcAMP concentrations. **(B)** Representative fluorescence-time traces for fcAMP binding to HCN1 reconstituted into POPC-POPE lipid. Traces are shown for 100 nM, 250 nM, 500 nM, 750 nM fcAMP concentrations.

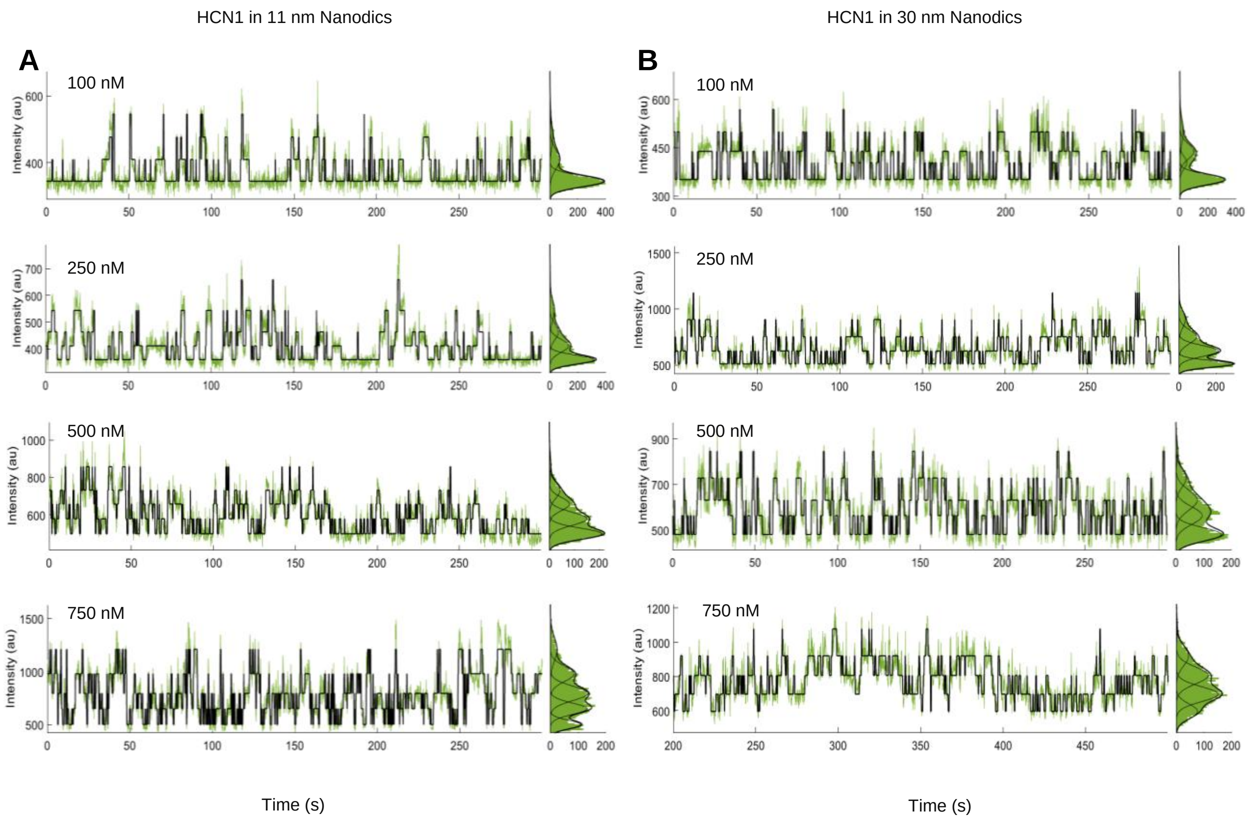

**Fig. S6. Representative Fluorescence-time traces for HCN1 in brain polar lipid spMSP1D1 (11nm) nanodiscs and HCN1 reconstituted in brain polar lipid spNW30 (30 nm) nanodiscs. (A)** Representative fluorescence-time traces for fcAMP binding to HCN1 in brain polar lipid spMSP1D1 (11 nm) nanodiscs overlaid with the idealized fit (black). Traces are shown for 100 nM, 250 nM, 500 nM, 750 nM fcAMP concentrations. **(B)** Representative fluorescence-time traces for fcAMP binding to HCN1 reconstituted in brain polar lipid spNW30 (30 nm) nanodiscs overlaid with the idealized fit (black). Traces are shown for 100 nM, 250 nM, 500 nM, 750 nM fcAMP concentrations.

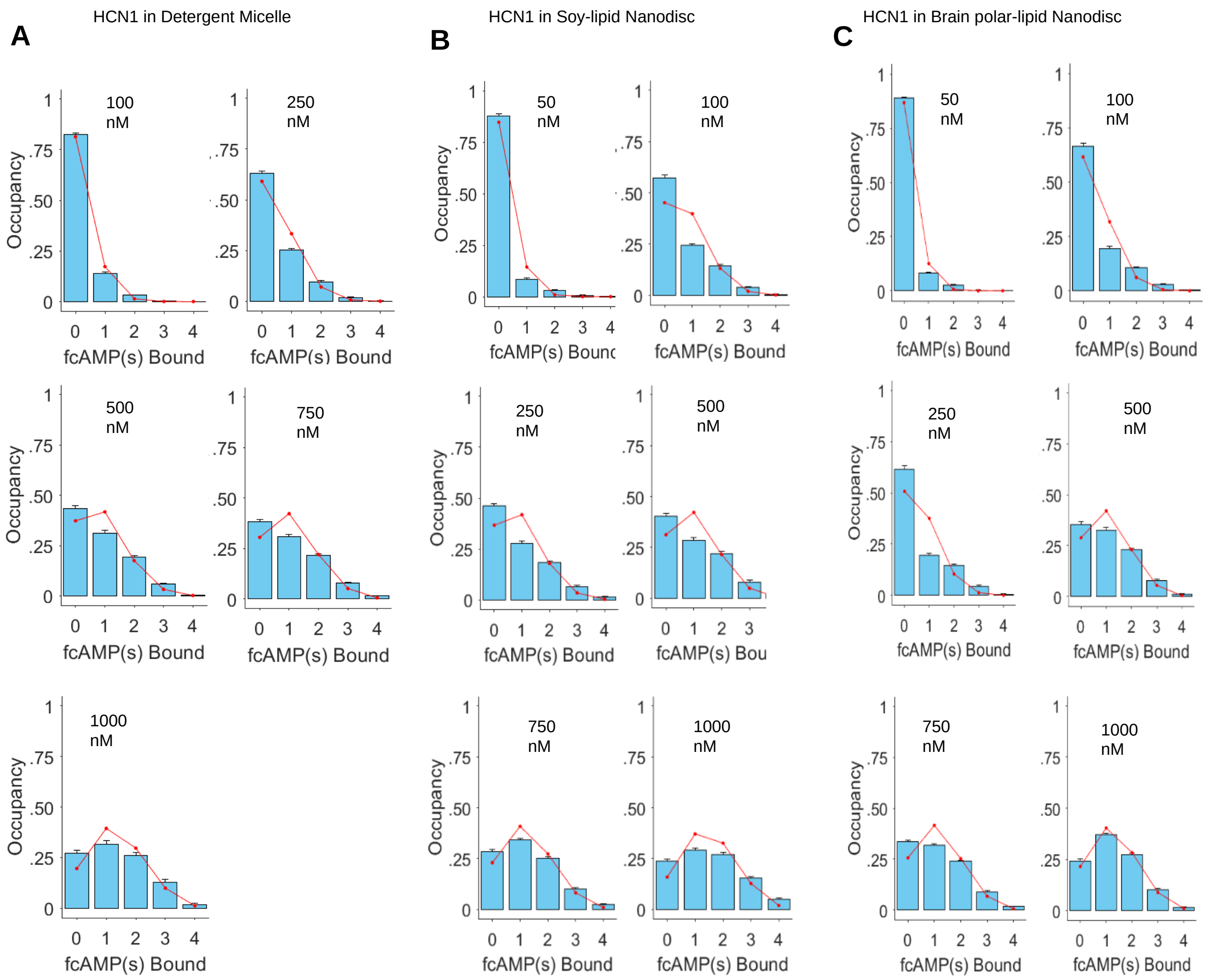

**Fig. S7. State occupancies for HCN1 in detergent micelles, soy polar lipid, and brain polar lipid nanodiscs.** Normalized mean state occupancy for each liganded state across all molecules of (**A**) HCN1 in detergent micelles and **(B)** HCN1 in soy polar lipid nanodiscs and (**C**) HCN1 in brain polar lipid nanodiscs at increasing fcAMP concentrations (mean ± s.e.m), overlaid with maximum likelihood estimations of a binomial distribution (red). Each plot indicates the total number of molecules (*n*) and data points included in the analysis. *P* is the success rate of the optimized binomial distribution considering four binding sites. All observed and expected state occupancy values are given in tables S10, S11 and S12.

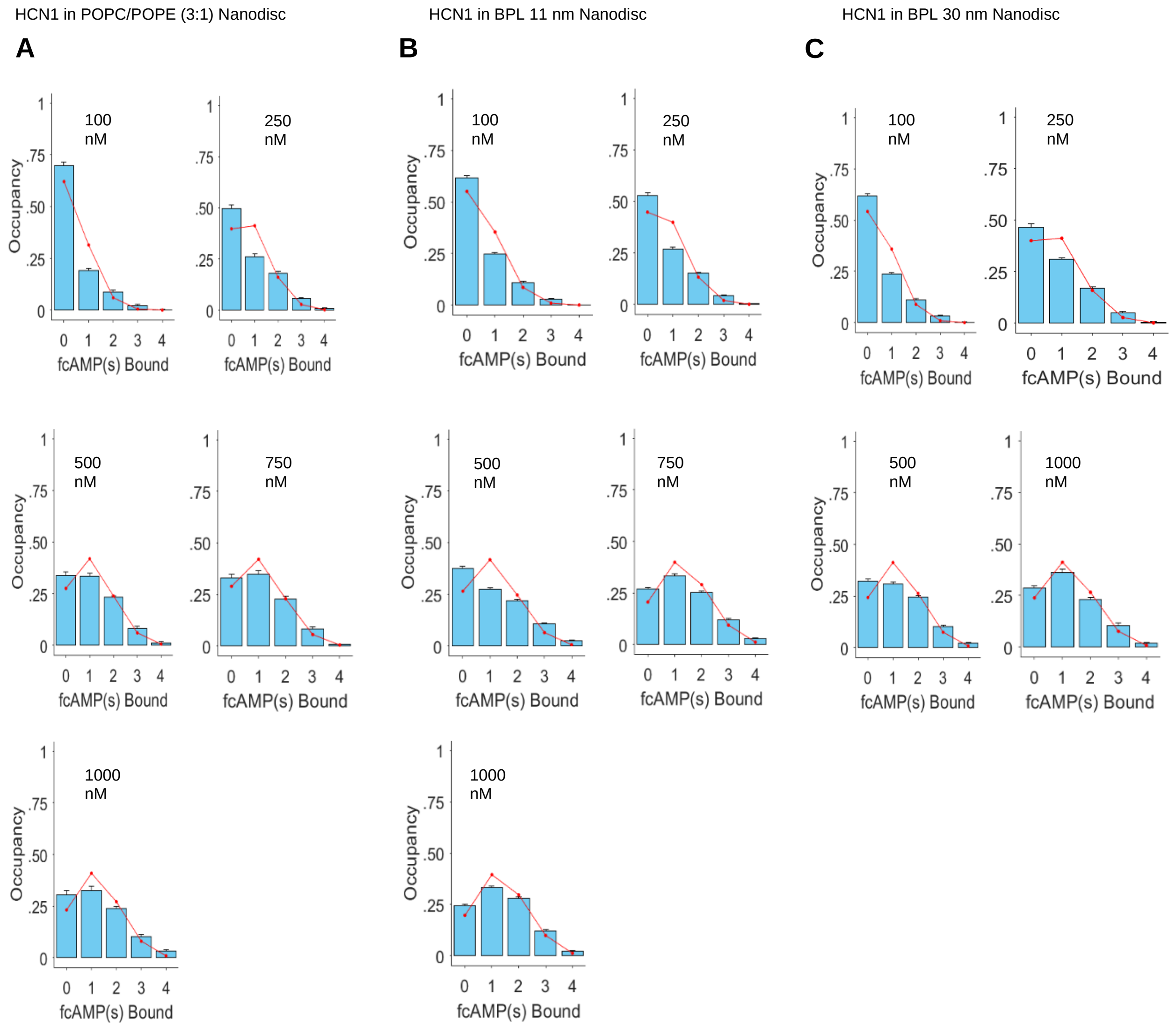

**Fig. S8. State occupancies for HCN1 reconstituted in POPC-POPE nanodiscs, brain polar lipid spMSP1D1 (11nm) nanodiscs, and HCN1 reconstituted in brain polar lipid spNW30 (30nm) nanodiscs.** Normalized mean state occupancy for each liganded state across all molecules of (**A**) HCN1 in POPC-POPE lipid nanodiscs **(B)** HCN1 in brain polar lipid spMSP1D1 (11nm) nanodiscs and **(C)** HCN1 reconstituted in brain polar lipid spNW30 (30 nm) nanodiscs at increasing fcAMP concentrations (mean ± s.e.m), overlaid with maximum likelihood estimations of a binomial distribution (red). Each plot indicates the total number of molecules (*n*) and data points included in the analysis. *P* is the success rate of the optimized binomial distribution considering four binding sites. All obtained and expected state occupancy values are in Supplementary Tables 13, 14 and 15.

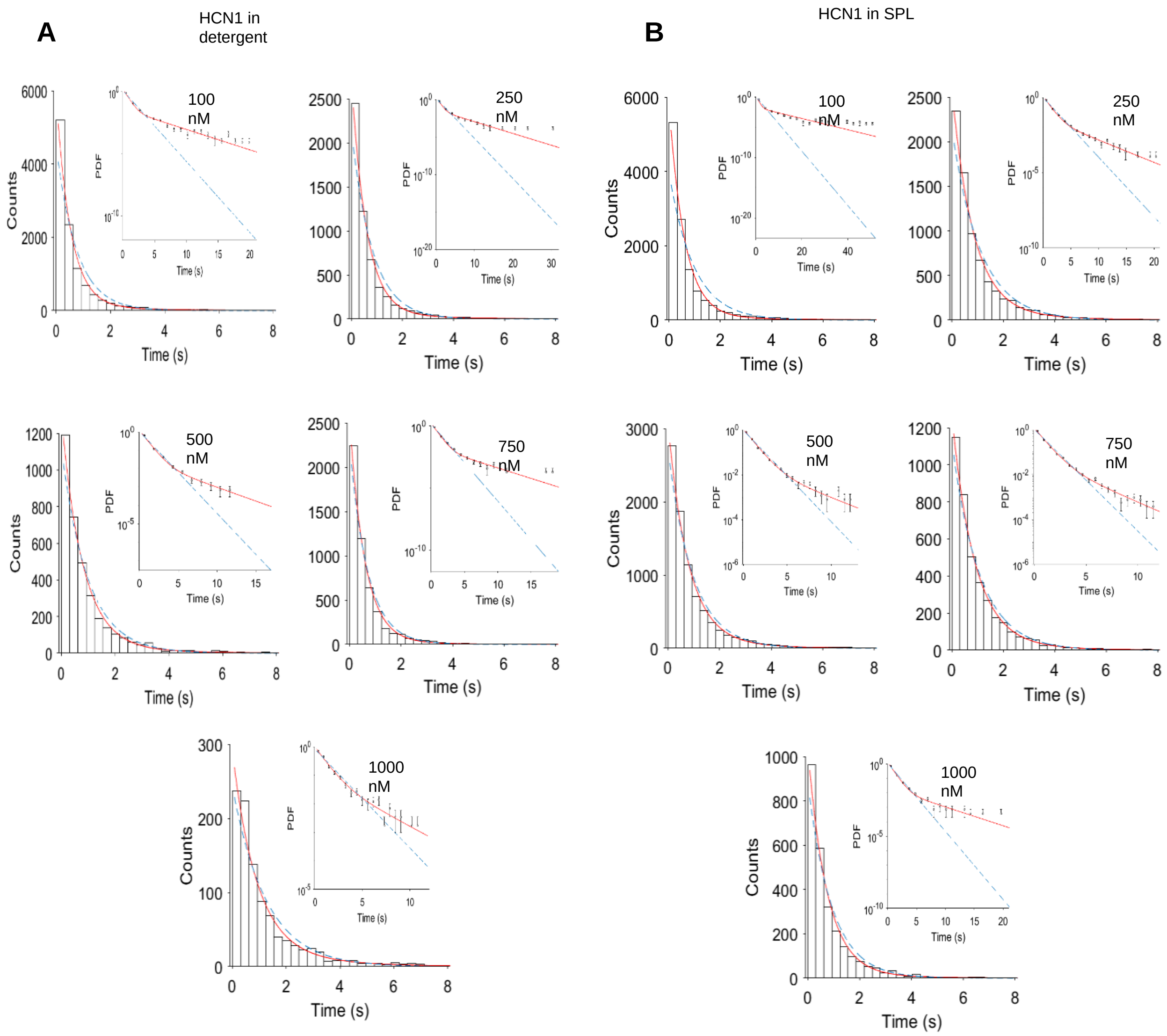

**Fig. S9. Isolated-B1 events of HCN1 in detergent (GDN) micelles and Soy Polar lipid nanodiscs. (A)** Dwell time distributions of isolated-B1 events for HCN1 in detergent micelle at various fcAMP concentrations. The distributions are overlaid with monoexponential (blue dashed) and bi-exponential (red) fits. Insets show probability density function of each distribution with logarithmic y-axis to highlights the population of the less frequent and long-lived states. Error bars correspond to the error in binomial distribution. The individual time constant values and associated amplitudes are given in the table S20. **(B)** Dwell time distributions of isolated-B1 events for HCN1 in soy polar lipid nanodiscs at various fcAMP concentrations. The distributions are overlaid with monoexponential (blue dashed) and bi-exponential (red) fits. Insets show probability density function of each distribution with logarithmic y-axis to highlights the population of the less frequent and long-lived states. Error bars are the same as shown in **a**. The individual time constant values and associated amplitudes are in table S19.

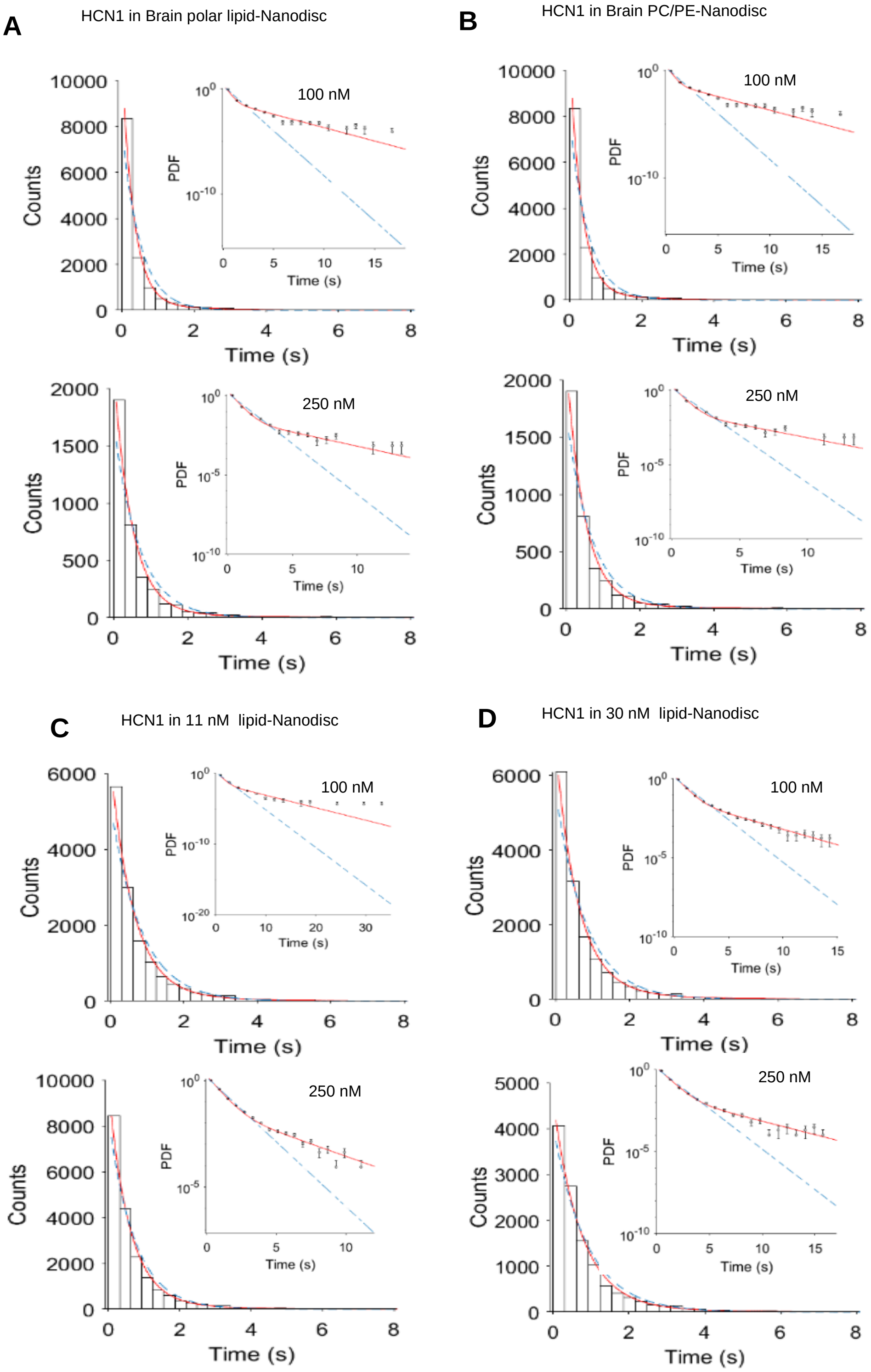

**Fig. S10. Isolated-B1 Events of HCN1 in Brain Polar lipid, spMSP1D1, spNW30 nanodiscs and POPC-POPE nanodiscs.** Dwell time distributions of isolated-B1 events for HCN1 in **(A)** Brain Polar lipid, **(B)** spMSP1D1 **(C)** POPC-POPE and **(D)** spNW30 nanodiscs at various fcAMP concentrations. The distributions are overlaid with monoexponential (blue dashed) and bi-exponential (red) fits. Insets show probability density function of each distribution with logarithmic y-axis to highlights the population of the less frequent and long-lived states. Error bars correspond to the error in binomial distribution. The individual time constant values and associated amplitudes are given in tables S21, S22, S23 and S24.

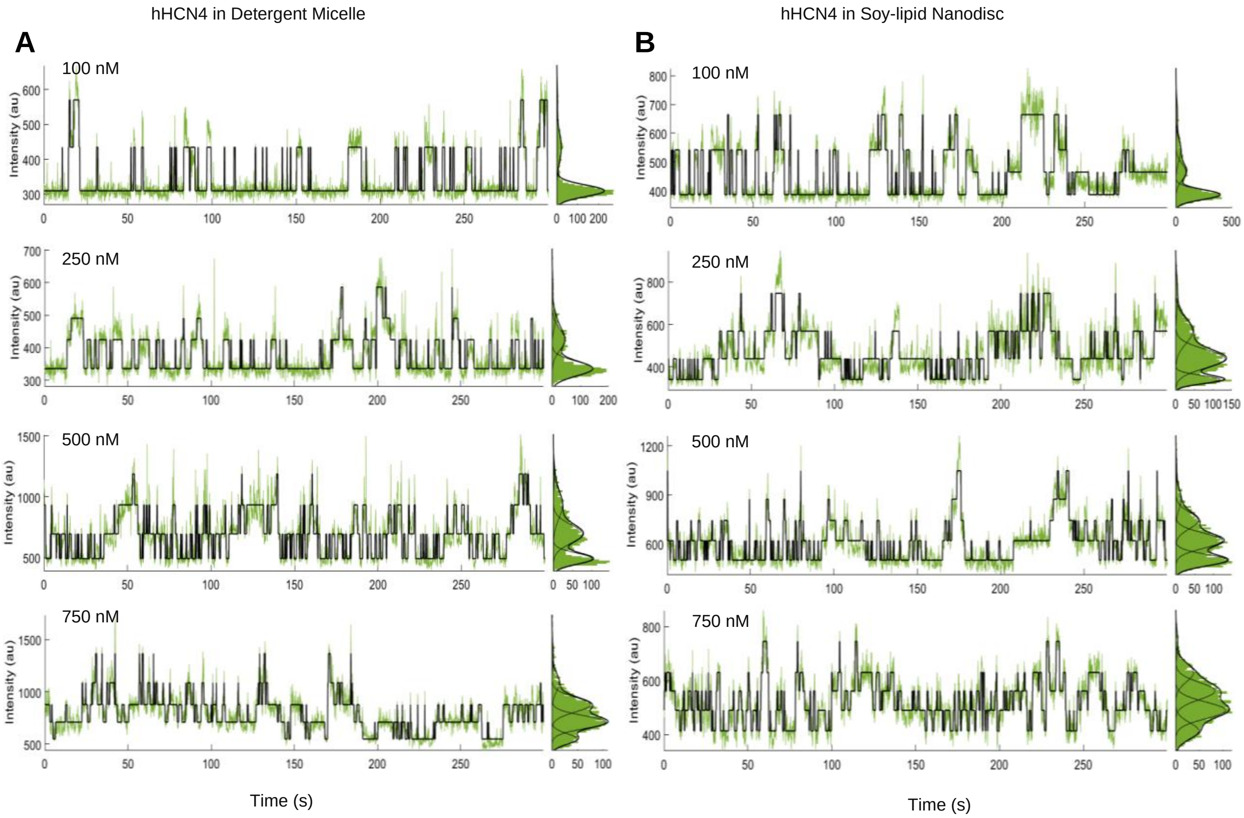

**Fig. S11. Representative Fluorescence-time traces for HCN4 in detergent micelles and HCN1 reconstituted in soy polar lipid nanodiscs. (A)** Representative fluorescence-time traces for fcAMP binding to HCN4 in detergent micelles overlaid with the idealized fit (black). Traces are shown for 100 nM, 250 nM, 500 nM, 750 nM fcAMP concentrations. **(B)** Representative fluorescence-time traces for fcAMP binding to HCN4 reconstituted in soy polar lipid nanodiscs overlaid with the idealized fit (black). Traces are shown for 100 nM, 250 nM, 500 nM, 750 nM fcAMP concentrations.

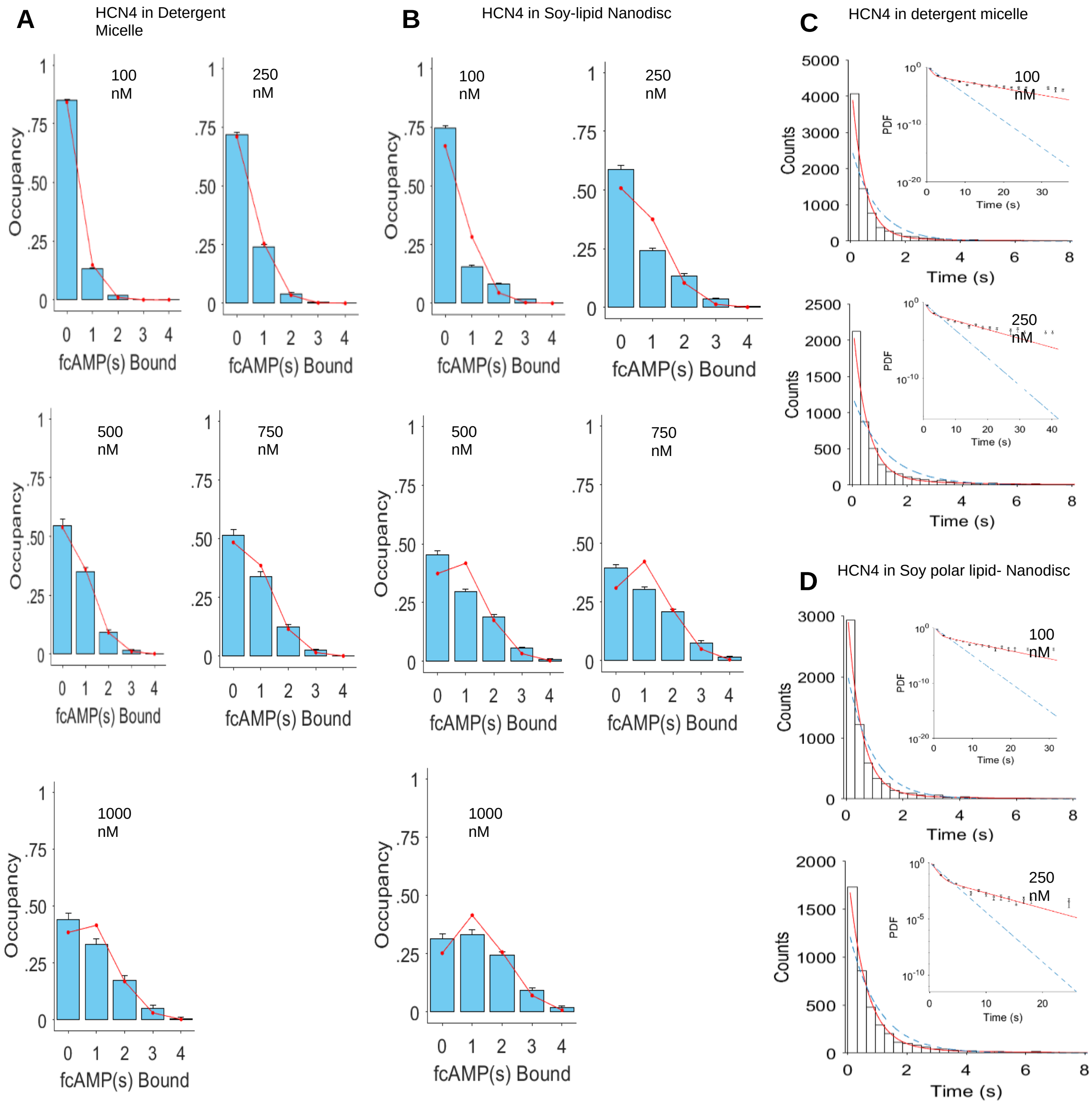

**Fig. S12. State occupancy and dwell time distributions of isolated-B1 events of HCN4 in detergent and soy polar lipid nanodiscs.** Normalized mean state occupancy for each liganded state across all molecules of (**A**) HCN4 in detergent micelle and (**B**) HCN4 in soy polar lipid nanodiscs at increasing fcAMP concentrations (mean ± s.e.m), overlaid with maximum likelihood estimations of a binomial distribution (red). Each plot indicates the total number of molecules (*n*) and data points included in the analysis. *P* is the success rate of the optimized binomial distribution considering four binding sites. All obtained and expected state occupancy values are in Supplementary Tables 16 and 17. Dwell time distributions of isolated-B1 events for (**C**) HCN4 in detergent micelle and (**D**) HCN4 reconstituted in soy polar nanodiscs at various fcAMP concentrations. The distributions are overlaid with monoexponential (blue dashed) and bi-exponential (red) fits. Insets show probability density function of each distribution with logarithmic y-axis to highlights the population of the less frequent and long-lived states. Error bars correspond to the error in binomial distribution. The individual time constant values and associated amplitudes are given in supplementary tables 26 and 27.

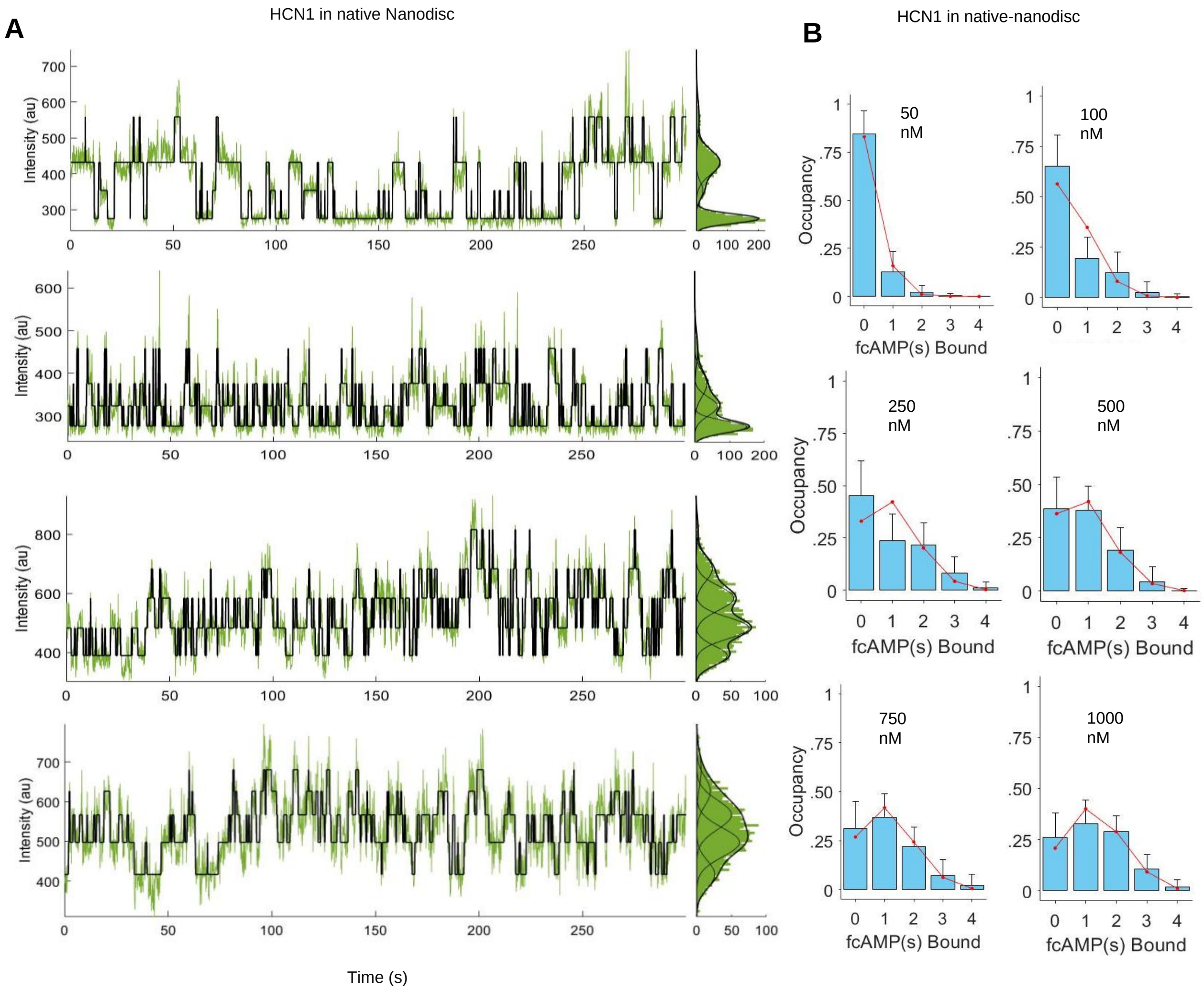

**Fig. S13. Representative fluorescence-time traces for HCN1 in native HEK membrane lipid nanodiscs. (A)** Representative fluorescence-time traces for fcAMP binding to HCN1 in native HEK membrane lipid nanodiscs overlaid with the idealized fit (black). Traces are shown for 100 nM, 250 nM, 500 nM, 750 nM fcAMP concentrations. **(B)** Normalized mean state occupancies for each liganded state across all molecules of HCN1 in native HEK membrane lipid nanodiscs at increasing fcAMP concentrations (mean ± s.e.m), overlaid with maximum likelihood estimations of a binomial distribution (red).

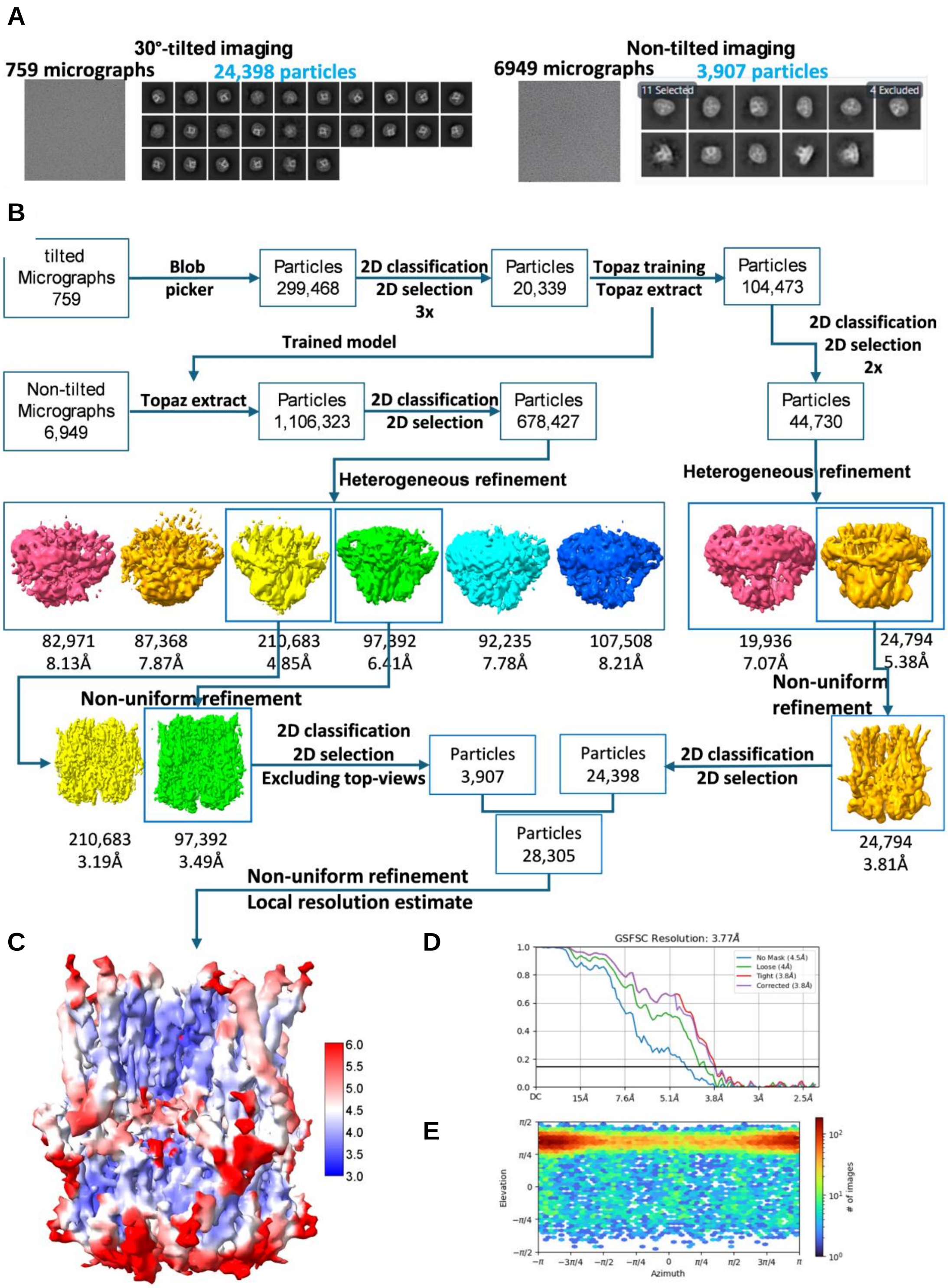

**Fig. S14. Cryo-EM data processing of hHCN1 in circular nanodisc.** (**A**) Examples of micrographs of tilted and non-tilted cryoEM imaging result with indicated number of curated micrographs and final 2D classes for particles used for final map refinement. **(B)** Flowchart for single particle analysis of two data sets (tilted and non-tilted), **(C)** The final map of hHCN1 in nanodisc with local resolution. **(D)** global resolution, **(E)** particle orientation distribution.

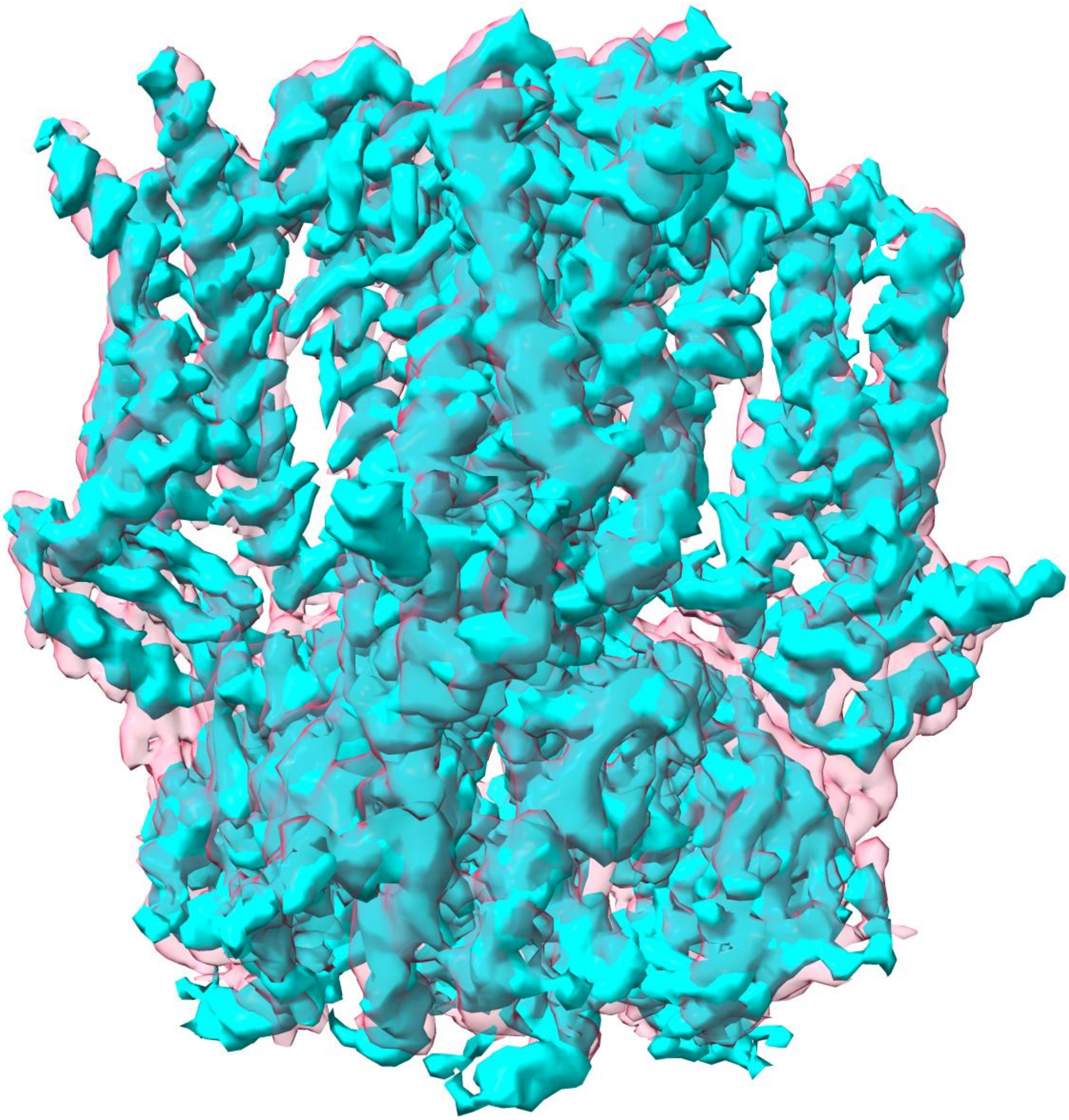

**Fig. S15. Comparison of hHCN1-nanodisc map (pink transparent) and hHCN1-detergent map (emd_8511, cyan).** Note that at similar contour level, except for unresolved end helix in the bottom, overall length of hHCN1-nanodisc is nearly identical to that in hHCN1-detergent.

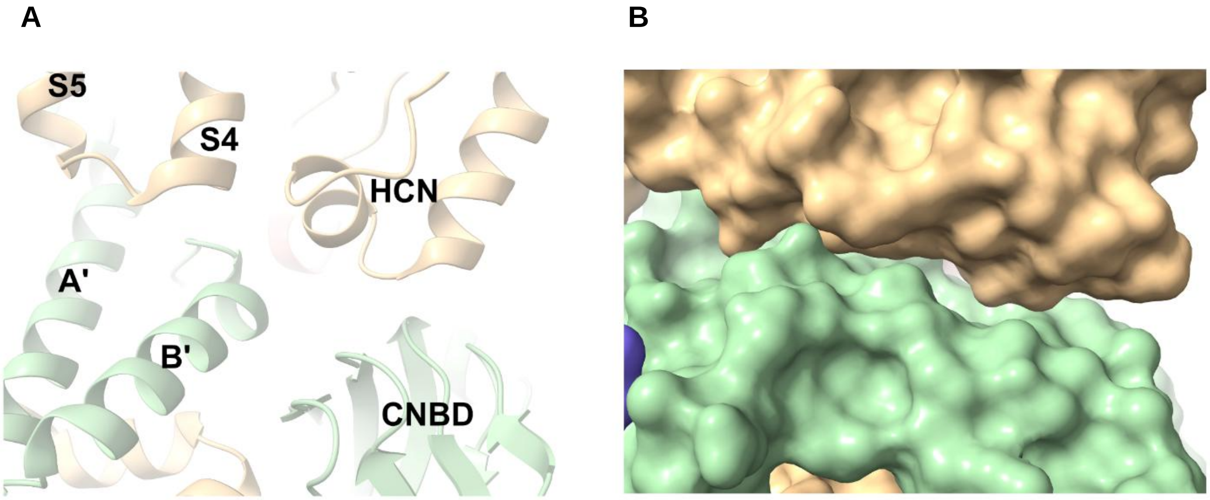

**Fig. S16. Subunit interface of hHCN1-nanodisc (8uc8) with ribbon (A) or surface (B) presentation.**

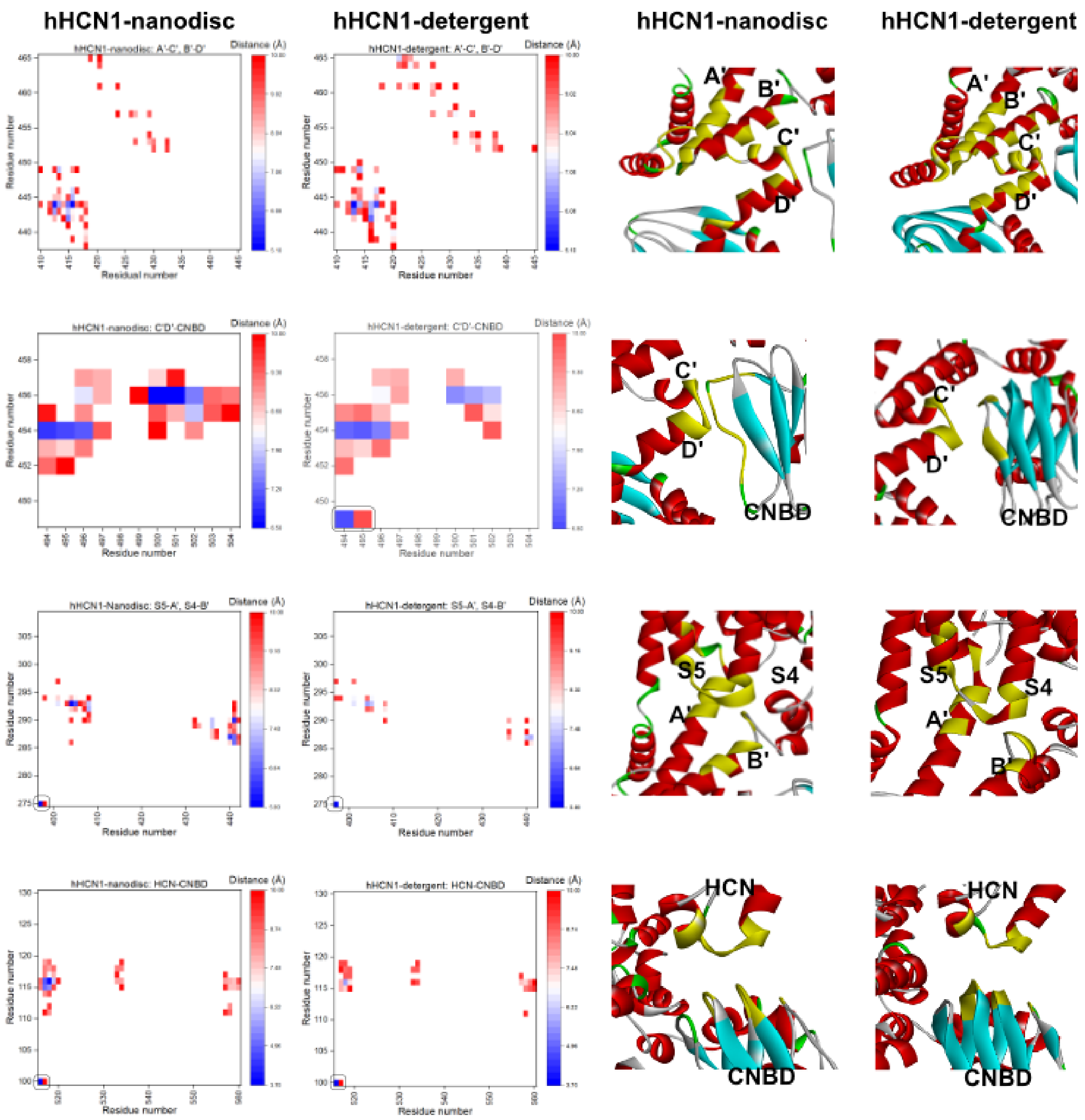

**Fig. S17. Detailed comparison of 2D contact maps in nanodisc and in detergent.** Note that left bottom dots are artificially added for matching minimum and maximum of each pair of figures. The residue-residue distances for the concatenated dimer of hHCN1-nanodisc or hHCN1-detergent (PDB: 5u6o) were calculated with UCSF Chimera 1.17.3. The distance matrices were exported to CSV files, which were then opened and processed in Excel with IFS() function to identify the contacting residues with cutoff of 10Å of the RR-distance. The 2D-contact map for all intersubunit residues identified is shown in Figure 4D. This figure shows important clusters of the 2D contact map in the subunit interface and their corresponding positions in the 3D structure as highlighted (yellow) and identified in the Discovery Studio Visualizer 2024 with the same cutoff.

**Table S1.**

**Summary of HCN1 in soy polar lipid nanodisc single molecule data**

| fcAMP concentration (nM) | Number of Molecules | Total Time (Minutes) | Total Events |
|---|---|---|---|
| 10 | 72 | 720 | 2105 |
| 50 | 60 | 600 | 6793 |
| 100 | 233 | 1396.68 | 47412 |
| 250 | 195 | 1237.93 | 47040 |
| 500 | 106 | 754 | 29648 |
| 750 | 171 | 1392 | 69776 |
| 1000 | 108 | 634.12 | 32085 |

**Table S2.**

**Summary of HCN1 in detergent micelles single molecule data**

| fcAMP concentration (nM) | Number of molecules | Total Time (Minutes) | Total Events |
|---|---|---|---|
| 50 | 183 | 915 | 12806 |
| 100 | 203 | 1190 | 22855 |
| 250 | 91 | 472 | 16311 |
| 500 | 53 | 278 | 11437 |
| 750 | 162 | 648 | 30569 |
| 1000 | 123 | 647 | 26777 |

**Table S3.**

**Summary of HCN1 in brain polar lipid nanodisc single molecule data**

| fcAMP concentration (nM) | Number of Molecules | Total Time (Minutes) | Total Events |
|---|---|---|---|

| | | | |
|---|---|---|---|
| 50 | 107 | 775 | 17129 |
| 100 | 173 | 1155 | 53271 |
| 250 | 59 | 440 | 20618 |
| 500 | 85 | 425 | 19956 |
| 750 | 278 | 1639.83 | 94156 |
| 1000 | 196 | 980 | 33150 |

**Table S4.**

**Summary of HCN1 in POPC-POPE nanodisc single molecule data**

| fcAMP concentration (nM) | Number of Molecules | Total Time (Minutes) | Total Events |
|---|---|---|---|
| 50 | 80 | 652.20 | 10383 |
| 100 | 60 | 552.46 | 21394 |
| 250 | 85 | 799.36 | 46022 |
| 500 | 58 | 557.81 | 30998 |
| 750 | 42 | 280.73 | 17558 |
| 1000 | 47 | 235 | 15245 |

**Table S5.**

**Summary of HCN1 in brain lipid 11nm nanodisc single molecule data**

| fcAMP concentration (nM) | Number of Molecules | Total Time (Minutes) | Total Events |
|---|---|---|---|
| 100 | 143 | 1424.66 | 54851 |
| 250 | 174 | 1739.64 | 87019 |
| 500 | 153 | 1530 | 84845 |
| 750 | 119 | 1190 | 72095 |
| 1000 | 154 | 1455 | 80593 |

**Table S6.**

**Summary of HCN1 in brain lipid 30nm nanodisc single molecule data**

| fcAMP concentration (nM) | Number of molecules | Total time (Minutes) | Total Events |
|---|---|---|---|
| 100 | 159 | 1577.54 | 61363 |
| 250 | 150 | 1491.1 | 59263 |
| 500 | 106 | 1060 | 58611 |
| 750 | 46 | 460 | 19697 |

**Table S7.**

**Summary of HCN4 in detergent micelles single molecule data**

| fcAMP concentration (nM) | Number of Molecules | Total Time (Minutes) | Total Events |
|---|---|---|---|
| 100 | 251 | 1255 | 16334 |
| 250 | 136 | 680 | 11128 |
| 500 | 52 | 260 | 5946 |
| 750 | 44 | 220 | 5753 |
| 1000 | 34 | 170 | 4821 |

**Table S8.**

**Summary of HCN4 in soy polar lipid nanodisc single molecule data**

| fcAMP concentration (nM) | Number of Molecules | Total Time (Minutes) | Total Events |
|---|---|---|---|
| 100 | 158 | 1105.98 | 24317 |
| 250 | 102 | 714 | 18796 |
| 500 | 89 | 623 | 19488 |
| 750 | 106 | 742 | 24360 |

| | | | |
|---|---|---|---|
| 1000 | 43 | 301 | 10468 |

**Table S9.**

**Summary of HCN1 in Native HEK membrane lipid nanodisc single molecule data**

| fcAMP concentration (nM) | Number of Molecules | Total Time (Minutes) | Total Events |
|---|---|---|---|
| 50 | 52 | 260 | 2279 |
| 100 | 67 | 335 | 10864 |
| 250 | 91 | 455 | 22456 |
| 500 | 146 | 730 | 26136 |
| 750 | 203 | 1015 | 48180 |
| 1000 | 112 | 560 | 30645 |

**Table S10.**

**Observed and expected state occupancies of HCN1 in soy polar lipid nanodiscs**

| OBSERVED | | | | | |
|---|---|---|---|---|---|
| | **Unbound** | **B1** | **B2** | **B3** | **B4** |
| **10 nM** | 0.928756944 | 0.056256944 | 0.01487963 | 0.000106481 | 0 |
| **50 nM** | 0.878555556 | 0.085638889 | 0.030075 | 0.005647222 | 8.33E-05 |
| **100 nM** | 0.573211135 | 0.242327709 | 0.143420937 | 0.037109593 | 0.003930626 |
| **250 nM** | 0.461810364 | 0.276024254 | 0.184210356 | 0.064265731 | 0.013689295 |
| **500 nM** | 0.401696541 | 0.285070755 | 0.218159591 | 0.07852044 | 0.016552673 |
| **750 nM** | 0.284879142 | 0.34013499 | 0.250614035 | 0.098715887 | 0.025655945 |
| **1000 nM** | 0.236339074 | 0.290212923 | 0.270923515 | 0.152106289 | 0.0504182 |
| **EXPECTED** | **Unbound** | **B1** | **B2** | **B3** | **B4** |
| **10 nM** | 0.91641944 | 0.080865152 | 0.002675838 | 3.94E-05 | 2.17E-07 |
| **50 nM** | 0.846638464 | 0.143923967 | 0.009174861 | 0.000259946 | 2.76E-06 |
| **100 nM** | 0.451587086 | 0.397167242 | 0.130989534 | 0.019200709 | 0.001055429 |
| **250 nM** | 0.366493535 | 0.418150937 | 0.178908551 | 0.034020963 | 0.002426014 |
| **500 nM** | 0.311889757 | 0.421842572 | 0.213959201 | 0.048231305 | 0.004077165 |
| **750 nM** | 0.229437935 | 0.408295238 | 0.272467479 | 0.080811378 | 0.00898797 |
| **1000 nM** | 0.158543438 | 0.370838127 | 0.3252758 | 0.126805068 | 0.018537567 |

**Table S11.**

**Observed and Expected State occupancies of HCN1 in detergent micelles**

| | | | | | |
|---|---|---|---|---|---|
| **OBSERVED** | | | | | |
| | **Unbound** | **B1** | **B2** | **B3** | **B4** |
| **50nM** | 0.895 | 0.099 | 0.005 | 0 | 0 |
| **100 nM** | 0.838 | 0.147 | 0.013 | 0.000027 | 0 |
| **250 nM** | 0.645 | 0.280 | 0.069 | 0.0045 | 0 |
| **500 nM** | 0.486 | 0.355 | 0.137 | 0.0206 | 0.000727 |
| **750 nM** | 0.441 | 0.334 | 0.179 | 0.0415 | 0.002721 |
| **1000 nM** | 0.359 | 0.395 | 0.200 | 0.0413 | 0.00246 |
| **EXPECTED** | | | | | |
| | **Unbound** | **B1** | **B2** | **B3** | **B4** |
| **50nM** | 0.8944 | 0.101 | 0.004 | 0.000008 | 5.7E-07 |
| **100 nM** | 0.846 | 0.144 | 0.009 | 0.000262 | 2.79E-06 |
| **250 nM** | 0.638 | 0.303 | 0.053 | 0.0042 | 0.000126 |
| **500 nM** | 0.503 | 0.376 | 0.105 | 0.0131 | 0.000615 |
| **750 nM** | 0.395 | 0.412 | 0.161 | 0.0281 | 0.00184 |
| **1000 nM** | 0.350 | 0.420 | 0.188 | 0.037 | 0.0028 |

**Table S12.**

**Observed and Expected State occupancies of HCN1 in brain polar lipid nanodiscs**

| | | | | | |
|---|---|---|---|---|---|
| **OBSERVED** | | | | | |
| | **Unbound** | **B1** | **B2** | **B3** | **B4** |
| **50** | 0.877861371 | 0.081655763 | 0.035990654 | 0.004492212 | 0 |
| **100** | 0.643926782 | 0.208594412 | 0.11477553 | 0.029815029 | 0.002888247 |

| | | | | | |
|---|---|---|---|---|---|
| **250** | 0.580649718 | 0.205344633 | 0.15930226 | 0.048878531 | 0.005824859 |
| **500** | 0.353356863 | 0.324133333 | 0.230533333 | 0.080352941 | 0.011623529 |
| **750** | 0.333383928 | 0.319302669 | 0.239366039 | 0.090208762 | 0.017738603 |
| **1000** | 0.236102041 | 0.372656463 | 0.275307823 | 0.101455782 | 0.014477891 |
| **EXPECTED** | **Unbound** | **B1** | **B2** | **B3** | **B4** |
| **50** | 0.849594447 | 0.141340831 | 0.008817691 | 0.000244489 | 2.54212E-06 |
| **100** | 0.59450145 | 0.33015645 | 0.068757159 | 0.006364049 | 0.000220892 |
| **250** | 0.473253906 | 0.389325979 | 0.120105758 | 0.016467655 | 0.000846703 |
| **500** | 0.286811774 | 0.420434006 | 0.231115975 | 0.056465032 | 0.005173214 |
| **750** | 0.257666303 | 0.415950611 | 0.25180084 | 0.067746999 | 0.006835247 |
| **1000** | 0.212074019 | 0.401748763 | 0.285399296 | 0.090109116 | 0.010668806 |

**Table S13.**

**Observed and Expected State occupancies of HCN1 in POPC-POPE lipid nanodiscs**

| **OBSERVED** | | | | | |
|---|---|---|---|---|---|
| | **Unbound** | **B1** | **B2** | **B3** | **B4** |
| **50** | 0.853421057 | 0.117053198 | 0.026038334 | 0.003487411 | 0 |
| **100** | 0.696747782 | 0.190961126 | 0.088135639 | 0.022614076 | 0.001541378 |
| **250** | 0.495554409 | 0.260161812 | 0.180455262 | 0.055861882 | 0.007966636 |
| **500** | 0.338681267 | 0.333462225 | 0.231330158 | 0.083459554 | 0.013066797 |
| **750** | 0.331099889 | 0.348746179 | 0.229253951 | 0.083039292 | 0.007860689 |
| **1000** | 0.305929078 | 0.325028369 | 0.237262411 | 0.100687943 | 0.031092199 |
| **EXPECTED** | | | | | |
| | **Unbound** | **B1** | **B2** | **B3** | **B4** |
| **50** | 0.83214025 | 0.156475958 | 0.011033924 | 0.000345804 | 4.06E-06 |
| **100** | 0.620798193 | 0.314324808 | 0.059681282 | 0.005036341 | 0.000159376 |
| **250** | 0.397561566 | 0.412441691 | 0.160454534 | 0.02774335 | 0.001798859 |
| **500** | 0.275784231 | 0.419116866 | 0.238853777 | 0.060498774 | 0.005746353 |
| **750** | 0.289571384 | 0.420694804 | 0.229197524 | 0.055497086 | 0.005039201 |
| **1000** | 0.231310211 | 0.408913562 | 0.271081258 | 0.079870233 | 0.008824735 |

**Table S14.**

**Observed and Expected State occupancies of HCN1 in brain lipid spMSP1D1 nanodiscs**

| OBSERVED | | | | | |
|---|---|---|---|---|---|
| | **Unbound** | **B1** | **B2** | **B3** | **B4** |
| 100 | 0.607071615 | 0.247623849 | 0.112579594 | 0.029573427 | 0.003152 |
| 250 | 0.528772992 | 0.268381361 | 0.150975607 | 0.044937089 | 0.006933 |
| 500 | 0.37514488 | 0.274238562 | 0.220367102 | 0.106339869 | 0.02391 |
| 750 | 0.268042017 | 0.332948179 | 0.251831933 | 0.119278711 | 0.027899 |
| 1000 | 0.244191558 | 0.332936147 | 0.281341991 | 0.120541126 | 0.020989 |
| **EXPECTED** | | | | | |
| | **Unbound** | **B1** | **B2** | **B3** | **B4** |
| 100 | 0.537722107 | 0.360873134 | 0.090820205 | 0.01015846 | 0.000426 |
| 250 | 0.445046261 | 0.399350013 | 0.134379654 | 0.020096981 | 0.001127 |
| 500 | 0.265161784 | 0.417416698 | 0.246410932 | 0.064649863 | 0.006361 |
| 750 | 0.205741091 | 0.398978136 | 0.290140545 | 0.093774602 | 0.011366 |
| 1000 | 0.19698949 | 0.394790206 | 0.296702327 | 0.099104418 | 0.012414 |

**Table S15.**

**Observed and Expected State occupancies of HCN1 in brain lipid spNW30 nanodiscs**

| **OBSERVED** | | | | | |
|---|---|---|---|---|---|
| | **Unbound** | **B1** | **B2** | **B3** | **B4** |
| 100 | 0.617856274 | 0.235744964 | 0.111013334 | 0.032603457 | 0.002781971 |
| 250 | 0.607754539 | 0.239916648 | 0.114683888 | 0.03412937 | 0.003515556 |
| 500 | 0.321647799 | 0.308633648 | 0.24604717 | 0.102628931 | 0.021042453 |
| 750 | 0.285405797 | 0.362456522 | 0.23173913 | 0.103192029 | 0.017206522 |
| **EXPECTED** | | | | | |
| | **Unbound** | **B1** | **B2** | **B3** | **B4** |
| 100 | 0.54204585 | 0.358710512 | 0.089019152 | 0.00981839 | 0.000406096 |
| 250 | 0.530323347 | 0.364502994 | 0.093949121 | 0.010762219 | 0.000462319 |
| 500 | 0.242584631 | 0.41229618 | 0.262776549 | 0.074435718 | 0.007906921 |
| 750 | 0.238616016 | 0.411171078 | 0.26569097 | 0.076304212 | 0.008217723 |

**Table S16.**

**Observed and Expected State occupancies of HCN4 in detergent micelles**

| **OBSERVED** | | | | | |
|---|---|---|---|---|---|
| | **Unbound** | **B1** | **B2** | **B3** | **B4** |
| **100** | 0.851671979 | 0.1314834 | 0.01649 | 0.000354582 | 0 |
| **250** | 0.721220588 | 0.239017157 | 0.037657 | 0.002041667 | 6.37E-05 |
| **500** | 0.549352564 | 0.346891026 | 0.089962 | 0.012923077 | 0.000872 |
| **750** | 0.516424242 | 0.33755303 | 0.120523 | 0.024613636 | 0.000886 |

| **1000** | 0.44427451 | 0.337 | 0.172353 | 0.044980392 | 0.001392 |
|---|---|---|---|---|---|
| **EXPECTED** | | | | | |
| | **Unbound** | **B1** | **B2** | **B3** | **B4** |
| **100** | 0.844466868 | 0.145816513 | 0.009442 | 0.000271728 | 2.93E-06 |
| **250** | 0.715838477 | 0.24958982 | 0.032634 | 0.001896403 | 4.13E-05 |
| **500** | 0.541261634 | 0.359105003 | 0.089344 | 0.009879385 | 0.00041 |
| **750** | 0.488464677 | 0.383282021 | 0.112781 | 0.014749219 | 0.000723 |
| **1000** | 0.398342913 | 0.412267563 | 0.160005 | 0.027599634 | 0.001785 |

**Table S17.**

**Observed and expected state occupancies of HCN4 in soy polar lipid nanodiscs**

| **OBSERVED** | | | | | |
|---|---|---|---|---|---|
| | **Unbound** | **B1** | **B2** | **B3** | **B4** |
| 100 | 0.74705004 | 0.154469 | 0.080171615 | 0.017304 | 0.001005959 |
| 250 | 0.587114846 | 0.241473 | 0.134568161 | 0.034456 | 0.002387955 |
| 500 | 0.452399679 | 0.294674 | 0.189026217 | 0.055265 | 0.008635634 |
| 750 | 0.395251572 | 0.302424 | 0.20916442 | 0.076388 | 0.016772237 |
| 1000 | 0.31406423 | 0.332896 | 0.243682171 | 0.091346 | 0.018012182 |
| **EXPECTED** | | | | | |
| | **Unbound** | **B1** | **B2** | **B3** | **B4** |
| 100 | 0.669731301 | 0.282395 | 0.044652536 | 0.003138 | 8.26969E-05 |
| 250 | 0.507705775 | 0.37503 | 0.103884489 | 0.012789 | 0.000590455 |
| 500 | 0.373453936 | 0.417083 | 0.174678757 | 0.032514 | 0.002269555 |
| 750 | 0.309293283 | 0.421794 | 0.215705973 | 0.049028 | 0.004178798 |
| 1000 | 0.251853387 | 0.414656 | 0.25601145 | 0.07025 | 0.007228838 |

**Table S18.**

**Observed and Expected State occupancies of HCN1 in HEK native membrane lipid nanodiscs**

| OBSERVED | | | | | |
|---|---|---|---|---|---|
| | **Unbound** | **B1** | **B2** | **B3** | **B4** |
| **50** | 0.845333333 | 0.129128205 | 0.022519230 | 0.003019230 | 0 |
| **100** | 0.651616915 | 0.195537313 | 0.123179104 | 0.025915422 | 0.003751243 |
| **250** | 0.451600732 | 0.237186813 | 0.214908424 | 0.083820512 | 0.012483516 |
| **500** | 0.385694064 | 0.377278539 | 0.192949772 | 0.042349315 | 0.00172831 |
| **750** | 0.312766831 | 0.370577997 | 0.222178982 | 0.073487685 | 0.020988506 |
| **1000** | 0.260089286 | 0.326574405 | 0.287666666 | 0.106836309 | 0.018833333 |
| **EXPECTED** | | | | | |
| | **Unbound** | **B1** | **B2** | **B3** | **B4** |
| **50** | 0.828984678 | 0.159181793 | 0.011462294 | .000366832 | .000004402 |
| **100** | 0.563313495 | 0.347639752 | 0.080452579 | 0.008274999 | .0003191738 |
| **250** | 0.329950031 | 0.421590177 | 0.202005904 | 0.043018486 | 0.003435401 |
| **500** | 0.362083284 | 0.418761572 | 0.181616973 | 0.035007696 | 0.002530474 |
| **750** | 0.268979667 | 0.418077832 | 0.2436834849 | 0.063126619 | 0.0061323967 |
| **1000** | 0.208286723 | 0.40011747 | 0.288233668 | 0.092282507 | 0.011079632 |

**Table S19.**

**Maximum likelihood estimation of isolated-B1 dwell times distributions of HCN1 in soy polar lipid nanodiscs**

| | Monoexponential Tau (sec) | | Biexponential Tau (sec) | | |
|---|---|---|---|---|---|
| **fcAMP concentration (nM)** | **τ(sec)** | **LL** | **τ(sec)** | **A** | **LL** |
| 10 | 2.9168±0.02 | -1430.7 | 0.7567±0.11<br>8.4116±1.57 | 0.7178±0.05<br>0.282±0.05 | -1190.6 |
| 50 | 1.03±0.04 | -2178.2 | 0.48±0.04<br>3.27±0.5 | 0.80±0.04<br>0.198±0.04 | -1806.9 |
| 100 | 0.964±0.02 | -12002 | 0.608±0.01<br>4.68±0.4 | 0.91±0.01<br>0.087±0.01 | -10147 |
| 250 | 1.1±0.06 | -8374.3 | 0.83±0.04<br>2.93±0.41 | 0.87±0.04<br>0.12±0.03 | -8124.6 |
| 500 | 1.068±0.03 | -4179.9 | 0.88±0.05<br>2.82±0.71 | 0.9±0.05<br>0.092±0.05 | -4113.7 |
| 750 | 0.95±0.02 | -8095.4 | 0.77±0.04<br>2.16±0.3 | 0.87±0.05<br>0.129±0.049 | -7976.3 |
| 1000 | 0.92±0.03 | -2422.5 | 0.74±0.05<br>3.37±1.03 | 0.93±0.24<br>0.06±0.03 | -2316.7 |

**Table S20.**

**Maximum likelihood estimation of isolated-B1 dwell times distributions of HCN1 in detergent micelles**

| | Monoexponential Tau (sec) | | Biexponential Tau (sec) | | |
|---|---|---|---|---|---|
| **fcAMP concentration (nM)** | **τ(sec)** | **LL** | **τ(sec)** | **A** | **LL** |
| 50 | 0.82±0.03 | -4531.8 | 0.54±0.02 | 0.89±0.02 | -3990 |

| | | | | | |
|---|---|---|---|---|---|
| | | | 3.12±0.44 | 0.10±0.02 | |
| 100 | 0.86±0.02 | -8433.5 | 0.59±0.02<br>3.14±0.34 | 0.89±0.02<br>0.10±0.02 | -7640.8 |
| 250 | 0.90±0.02 | -5029.5 | 0.62±0.03<br>3.2±0.5 | 0.89±0.03<br>0.10±0.02 | -4592.7 |
| 500 | 0.96±0.03 | -2836.6 | 0.72±0.05<br>2.96±0.68 | 0.89±0.05<br>0.1±0.04 | -2707.7 |
| 750 | 0.89±0.02 | -4530 | 0.75±0.04<br>3.32±0.84 | 0.94±0.03<br>0.05±0.03 | -4368.8 |
| 1000 | 1.27±0.04 | -5691.8 | 1.09±0.05<br>3.34±0.84 | 0.92±0.05<br>0.07±0.04 | -5567.2 |

**Table S21.**

**Maximum likelihood estimation of isolated-B1 dwell times distributions of HCN1 in brain polar lipid nanodiscs**

| | **Monoexponential Tau (sec)** | | **Biexponential Tau (sec)** | | |
|---|---|---|---|---|---|
| **fcAMP concentration (nM)** | **τ (sec)** | **LL** | **τ (sec)** | **A** | **LL** |
| 50 | 0.58±0.02 | -2487.7 | 0.4±0.02<br>2.26±0.31 | 0.9±0.02<br>0.09±0.02 | -2019.5 |
| 100 | 0.50±0.01 | -4214.7 | 0.33±0.01<br>1.70±0.14 | 0.87±0.02<br>0.125±0.01 | -3133.7 |
| 250 | 0.68±0.03 | -2333.2 | 0.49±0.03<br>2.4±0.4 | 0.90±0.03<br>0.095±0.03 | -2086.2 |
| 500 | 1.06±0.04 | -2274.2 | 0.95±0.06<br>2.81±1.24 | 0.94±0.06<br>0.057±0.05 | -2260 |
| 750 | 0.92±0.02 | -8531.6 | 0.79±0.03<br>2.3±0.5 | 0.91±0.03<br>0.08±0.04 | -8429.7 |
| 1000 | 1.76±0.07 | -3956.7 | 1.67±0.09 | 0.98±0.02 | -3938.4 |

| | | | | |
|---|---|---|---|---|
| | | | 7.3±4.6 | 0.016±0.019 |

**Table S22.**

**Maximum likelihood estimation of isolated-B1 dwell times distributions of HCN1 in POPC-POPE Lipid Nanodiscs**

| | Monoexponential Tau (sec) | | Biexponential Tau (sec) | | |
|---|---|---|---|---|---|
| **fcAMP Concentration (nM)** | **τ (sec)** | **LL** | **τ (sec)** | **A** | **LL** |
| 100 | 0.66±0.01 | -3858.6 | 0.53±0.03<br>1.76±0.28 | 0.88±0.03<br>0.11±0.04 | -3708.9 |
| 250 | 0.67±0.02 | -5106.2 | 0.54±0.02<br>2.18±0.33 | 0.92±0.02<br>0.07±0.02 | -4849.1 |
| 500 | 0.89±0.02 | -3654.7 | 0.78±0.11<br>2.58±0.75 | 0.93±0.04<br>0.06±0.04 | -3601.5 |
| 750 | 0.82±0.03 | -1862.7 | 0.65±0.05<br>2.58±0.6 | 0.90±0.04<br>0.09±0.05 | -1785.5 |
| 1000 | 0.70±0.03 | -1093.8 | 0.51±0.04<br>2.16±0.59 | 0.88±0.05<br>0.11±0.05 | -1015.2 |

**Table S23.**

**Maximum likelihood estimation of isolated-B1 dwell times distributions of HCN1 in spMSP1D1 (11nm Circular) Brain Lipid Nanodiscs**

| | Monoexponential Tau (sec) | | Biexponential Tau (sec) | | |
|---|---|---|---|---|---|
| **fcAMP Concentration (nM)** | **τ(sec)** | **LL** | **τ(sec)** | **A** | **LL** |
| 100 | 0.82±0.02 | -1100.9 | 0.62±0.02<br>2.47±0.28 | 0.89±0.02<br>0.104±0.02 | -10478 |
| 250 | 0.71±0.01 | -12935 | 0.57±0.01<br>1.88±0.19 | 0.89±0.02<br>0.106±0.02 | -12529 |
| 500 | 0.82±0.02 | -8157.3 | 0.69±0.03<br>1.98±0.35 | 0.90±0.04<br>0.09±0.04 | -8036.6 |
| 750 | 0.87±0.02 | -6109.5 | 0.76±0.04<br>2.3±0.64 | 0.93±0.21<br>0.06±0.03 | -6035.7 |
| 1000 | 1.02±0.03 | -6577 | 0.97±0.03<br>3.47±1.3 | 0.98±0.02<br>0.018±0.01 | -6553.7 |

**Table S24.**

**Maximum likelihood estimation of isolated-B1 dwell times distributions of HCN1 in spNW30 (30nm Circular) Brain Lipid Nanodiscs**

| | Monoexponential Tau (sec) | | Biexponential Tau (sec) | | |
|---|---|---|---|---|---|
| **fcAMP Concentration (nM)** | **τ(sec)** | **LL** | **τ(sec)** | **A** | **LL** |
| 100 | 0.80±0.1 | -11538 | 0.60±0.02<br>2.22±0.2 | 0.87±0.02<br>0.127±0.02 | -11004 |
| 250 | 0.88±0.02 | -10253 | 0.73±0.03<br>2.65±0.37 | 0.91±0.02<br>0.08±0.03 | -9977.3 |
| 500 | 0.92±0.05 | -5982.1 | 0.86±0.03 | 0.97±0.02 | -5930.6 |

| | | | | | |
|---|---|---|---|---|---|
| | | | 3.20±1.10 | 0.02±0.02 | |
| 750 | 1.49±0.08 | -1859.9 | 1.49±0.09 | 0.96±76.7 | -1859.9 |
| | | | 1.49±0.03 | 0.03±77.6 | |

**Table S25.**

**Maximum likelihood estimation of isolated-B1 dwell times distributions of HCN1 in native HEK membrane nanodiscs**

| | **Monoexponential Tau (sec)** | | **Biexponential Tau (sec)** | | |
|---|---|---|---|---|---|
| **fcAMP concentration (nM)** | **τ (sec)** | **LL** | **τ (sec)** | **A** | **LL** |
| 50 | 2.26±0.02 | -1151.6 | 1.02±0.02<br>8.02±0.43 | 0.822±0.02<br>0.177±0.02 | -1016.9 |
| 100 | 0.95±0.02 | -2397.5 | 0.71±0.02<br>3.36±0.43 | 0.91±0.02<br>0.087±0.02 | -2263.5 |
| 250 | 0.94±0.02 | -2531.4 | 0.78±0.03<br>2.19±0.32 | 0.88±0.04<br>0.12±0.04 | -2496.5 |
| 500 | 1.57±0.02 | -5923 | 1.35±0.03<br>5.55±0.78 | 0.95±0.01<br>0.05±0.01 | -58367 |
| 750 | 1.28±0.02 | -6817.4 | 1.23±0.02<br>3.38±0.78 | 0.98±0.01<br>0.02±0.01 | -6810.5 |
| 1000 | 0.95±0.02 | -2642 | 0.84±0.03<br>2.52±0.48 | 0.94±0.03<br>0.06±0.03 | -2621 |

**Table S26.**

**Maximum likelihood estimation of isolated-B1 dwell times distributions of HCN4 in Detergent Micelles**

| | **Monoexponential Tau (sec)** | | **Biexponential Tau (sec)** | | |
|---|---|---|---|---|---|
| **fcAMP Concentration (nM)** | **τ(sec)** | **LL** | **τ(sec)** | **A** | **LL** |
| 100 | 0.92±0.02 | -7230.5 | 0.47±0.01<br>3.70±0.4 | 0.86±0.02<br>0.14±0.02 | -5659.9 |
| 250 | 1.18±0.03 | -5622 | 0.51±0.03 | 0.78±0.03 | -4706.6 |

| | | | | | |
|---|---|---|---|---|---|
| | | | 3.63±0.34 | 0.21±0.03 | |
| 500 | 1.00±0.05 | -1847.8 | 0.52±0.21<br>2.76±0.49 | 0.78±0.05<br>0.21±0.07 | -1639.1 |
| 750 | 1.08±0.06 | -1410.7 | 0.6±0.07<br>3.5±0.83 | 0.83±0.06<br>0.16±0.06 | -1246 |
| 1000 | 1.106 | -985.4 | 0.55±0.07<br>3.2±0.74 | 0.79±0.06<br>0.20±0.06 | -868.2 |

**Table S27.**

**Maximum likelihood estimation of isolated-B1 dwell times distributions of HCN4 in Soy Polar Lipid Nanodiscs**

| | Monoexponential Tau (sec) | | Biexponential Tau (sec) | | |
|---|---|---|---|---|---|
| **fcAMP Concentration (nM)** | **τ(sec)** | **LL** | **τ(sec)** | **A** | **LL** |
| 100 | 0.85±0.02 | -4890.3 | 0.48±0.02<br>2.97±0.31 | 0.85±0.02<br>0.14±0.02 | -4122.4 |
| 250 | 0.98±0.04 | -3924.6 | 0.59±0.03<br>3.19±0.44 | 0.85±0.03<br>0.14±0.03 | -3516.6 |
| 500 | 1.03±0.03 | -3347.1 | 0.67±0.05<br>2.83±0.45 | 0.67±0.05<br>0.16±0.04 | -3143.4 |
| 750 | 1.09±0.04 | -3525.7 | 0.76±0.06<br>2.82±0.50 | 0.83±0.05<br>0.16±0.05 | -3384.5 |
| 1000 | 1.44±0.09 | -1246.7 | 1.19±0.23<br>3.15±0.82 | 0.87±0.21<br>0.12±0.09 | -1235.7 |

**Table S28.**

**Optimized Rate constants for Sequential binding model**

| | **Model** | **Parameter** | **Rate** | **s.e.m.** |
|---|---|---|---|---|
| | | *k*U,B1 | 3.98E+05 | 5.85E+03 |
| | | *k*B1,U | 1.02E+00 | 3.78E-03 |
| | | *k*B1,B2 | 3.98E+05 | 5.15E+03 |
| **HCN1 in BPL Nanodiscs** | 1 | *k*B2,B1 | 5.40E-01 | 4.62E-03 |
| | | *k*B2,B3 | 3.01E+05 | 4.24E+03 |
| | | *k*B3,B2 | 3.78E-01 | 7.74E-03 |
| | | *k*B3,B4 | 3.01E+05 | 4.82E+03 |
| | | *k*B4,B3 | 3.18E-01 | 2.01E-02 |

**Table S29.**

**Optimized Rate constants for Sequential binding model**

| | **Model** | **Parameter** | **Rate** | **s.e.m.** |
|---|---|---|---|---|
| | | *k*U,B1 | 4.06E+05 | 7.60E+03 |
| | | *k*B1,U | 8.41E-01 | 3.96E+00 |
| | | *k*B1,B2 | 4.60E+05 | 7.60E+03 |
| **HCN1 in SPL Nanodiscs** | 1 | *k*B2,B1 | 4.81E-01 | 5.31E+00 |
| | | *k*B2,B3 | 3.88E+05 | 6.89E+03 |
| | | *k*B3,B2 | 3.32E-01 | 8.57E+00 |
| | | *k*B3,B4 | 3.61E+05 | 6.90E+03 |
| | | *k*B4,B3 | 2.44E-01 | 1.74E+01 |

**Table S30.**

**Optimized Rate constants for Sequential binding model**

| | Model | Parameter | Rate | s.e.m. |
|---|---|---|---|---|
| | | *k*U,B1 | 5.49E+05 | 1.04E+04 |
| | | *k*B1,U | 1.18E+00 | 5.70E-03 |
| | | *k*B1,B2 | 5.58E+05 | 9.80E+03 |
| **HCN1 in PCPE Nanodiscs** | 1 | *k*B2,B1 | 5.96E-01 | 7.10E-03 |
| | | *k*B2,B3 | 4.71E+05 | 9.40E+03 |
| | | *k*B3,B2 | 4.39E-01 | 1.26E-02 |
| | | *k*B3,B4 | 4.52E+05 | 1.04E+04 |
| | | *k*B4,B3 | 3.63E-01 | 3.31E-02 |

**Table S31.**

**Optimized Rate constants for Sequential binding model**

| | Model | Parameter | Rate | s.e.m. |
|---|---|---|---|---|
| | | *k*U,B1 | 3.28E+05 | 8.10E+03 |
| | | *k*B1,U | 9.75E-01 | 6.00E-03 |
| | | *k*B1,B2 | 2.88E+05 | 8.20E+03 |
| **HCN1 in detergent micelle** | 1 | *k*B2,B1 | 4.93E-01 | 9.30E-03 |
| | | *k*B2,B3 | 2.09E+05 | 7.20E+03 |
| | | *k*B3,B2 | 3.75E-01 | 2.03E-02 |
| | | *k*B3,B4 | 1.68E+05 | 9.50E+03 |
| | | *k*B4,B3 | 2.83E-01 | 6.23E-02 |

**Table S32.**

**Optimized Rate constants for Sequential binding model**

| | Model | Parameter | Rate | s.e.m. |
|---|---|---|---|---|
| | | *k*U,B1 | 4.58E+05 | 5.40E+03 |

| | | | | |
|---|---|---|---|---|
| | | *k*B1,U | 1.01E+00 | 2.90E-03 |
| | | *k*B1,B2 | 4.56E+05 | 4.70E+03 |
| **HCN1 in 11 nm (spMSP1D1) Circular Nanodiscs** | 1 | *k*B2,B1 | 5.40E-01 | 3.70E-03 |
| | | *k*B2,B3 | 3.98E+05 | 4.00E+03 |
| | | *k*B3,B2 | 3.98E-01 | 6.10E-03 |
| | | *k*B3,B4 | 3.57E+05 | 4.10E+03 |
| | | *k*B4,B3 | 3.16E-01 | 1.49E-02 |

**Table S33.**

**Optimized Rate constants for Sequential binding model**

| | **Model** | **Parameter** | **Rate** | **s.e.m.** |
|---|---|---|---|---|
| | | *k*U,B1 | 5.83E+05 | 9.20E+03 |
| | | *k*B1,U | 9.31E-01 | 3.60E-03 |
| | | *k*B1,B2 | 6.26E+05 | 9.00E+03 |
| **HCN1 in 30 nm Circular (spNW30) Nanodisc** | 1 | *k*B2,B1 | 5.33E-01 | 5.10E-03 |
| | | *k*B2,B3 | 5.57E+05 | 8.60E+03 |
| | | *k*B3,B2 | 3.88E-01 | 8.80E-03 |
| | | *k*B3,B4 | 4.74E+05 | 8.90E+03 |
| | | *k*B4,B3 | 3.26E-01 | 2.41E-02 |

**Table S34.**

**Optimized Rate constants for Sequential binding model**

| | **Model** | **Parameter** | **Rate** | **s.e.m.** |
|---|---|---|---|---|
| | | *k*U,B1 | 1.28E+06 | 6.25E+03 |
| | | *k*B1,U | 7.26E-01 | 3.61E-03 |
| | | *k*B1,B2 | 9.99E+05 | 5.31E+03 |

| | | | | |
|---|---|---|---|---|
| **HCN1 in native HEK membrane Nanodiscs** | 1 | *k*B2,B1 | 9.62E-01 | 5.06E-03 |
| | | *k*B2,B3 | 5.63E+05 | 4.83E+03 |
| | | *k*B3,B2 | 1.14E+00 | 1.01E-02 |
| | | *k*B3,B4 | 3.12E+05 | 6.41E+03 |
| | | *k*B4,B3 | 1.12E+00 | 2.27E-02 |

**Table S35.**

**Optimized Rate constants for Sequential binding model**

| | **Model** | **Parameter** | **Rate** | **s.e.m.** |
|---|---|---|---|---|
| | | *k*U,B1 | 2.44E+05 | 5.30E+03 |
| | | *k*B1,U | 7.17E-01 | 3.90E-03 |
| | | *k*B1,B2 | 3.18E+05 | 6.50E+03 |
| **HCN4 in SPL Nanodisc** | 1 | *k*B2,B1 | 3.69E-01 | 5.00E-03 |
| | | *k*B2,B3 | 2.50E+05 | 5.80E+03 |
| | | *k*B3,B2 | 2.86E-01 | 9.80E-03 |
| | | *k*B3,B4 | 2.60E+05 | 7.10E+03 |
| | | *k*B4,B3 | 2.39E-01 | 2.63E-02 |

**Table S36.**

**Optimized Rate constants for Sequential binding model**

| | **Model** | **Parameter** | **Rate** | **s.e.m.** |
|---|---|---|---|---|
| | | *k*U,B1 | 2.20E+05 | 6.00E+03 |
| | | *k*B1,U | 7.22E-01 | 4.90E-03 |
| | | *k*B1,B2 | 1.85E+05 | 6.70E+03 |
| **HCN4 in detergent micelle** | 1 | *k*B2,B1 | 4.64E-01 | 1.22E-02 |
| | | *k*B2,B3 | 1.53E+05 | 8.80E+03 |
| | | *k*B3,B2 | 3.62E-01 | 3.16E-02 |

| | | | | |
|---|---|---|---|---|
| | | *k*B3,B4 | 1.21E+05 | 1.22E+04 |
| | | *k*B4,B3 | 2.90E-01 | 1.18E-01 |

**Movie S1.**
hHCN1 detergent to nanodisc morph top view (separate file).

**Movie S2.**
hHCN1 detergent to nanodisc morph side view (separate file).
